# Distinct circulating monocytes up-regulate CD52 and sustain innate immune function in patients with cirrhosis unless acute decompensation emerges

**DOI:** 10.1101/2024.04.03.587894

**Authors:** Anne Geng, Robert G. Brenig, Julien Roux, Mechthild Lütge, Hung-Wei Cheng, Emilio Flint, Marie-Anne Meier, Oltin T. Pop, Patrizia Künzler-Heule, Mark J. W. McPhail, Savas Soysal, David Semela, Markus Heim, Chris J. Weston, Burkhard Ludewig, Christine Bernsmeier

**Affiliations:** Department of Biomedicine, University of Basel, Switzerland; University Centre for Gastrointestinal and Liver Disease Basel, Switzerland; Bioinformatics Core Facility, Department of Biomedicine, University of Basel, Switzerland; Swiss Institute of Bioinformatics, Basel, Switzerland; Institute of Immunobiology, Cantonal Hospital St. Gallen, Switzerland; Liver Biology Laboratory, Division of Gastroenterology and Hepatology, Cantonal Hospital St. Gallen, Switzerland; Institute of Liver Studies, King’s College Hospital, and School of Immunology and Microbial Sciences, King’s College London, London, United Kingdom; Institute of Immunology and Immunotherapy, NIHR Biomedical Research Unit and Centre for Liver Research, The Medical School, University of Birmingham, Birmingham, United Kingdom

**Author notes:** **Grant support** Swiss National Science Foundation, Grant Nr: 320030_189072; Clarunis Research Funds. **Corresponding author contact information:** Prof. Christine Bernsmeier, MD, PhD, Department of Biomedicine, University of Basel, University Centre for Gastrointestinal and Liver Diseases Hebelstrasse 20, 4031 Basel, Switzerland.

**Keywords:** Cirrhosis, AD, ACLF, NAD, innate immunity, immuneparesis, scRNAseq, monocytes, macrophages, M-MDSC, CD52, PLC

## Abstract

**Background & Aims:** Infectious complications determine the prognosis of cirrhosis patients. Their infection susceptibility relates to the development of immuneparesis, a complex interplay of different immunosuppressive cells and soluble factors. Mechanisms underlying the dynamics of immuneparesis of innate immunity remain inconclusive. We aimed to dissect the heterogeneity of circulating monocyte states in different cirrhosis stages, and pursued the function of selected differentially expressed (DE) genes.

**Methods:** We systematically investigated circulating monocytes in health, compensated and not-acutely decompensated (NAD) cirrhosis using single cell RNA sequencing. Selective genes were confirmed by flow cytometry and diverse functional assays on monocytes *ex vivo*.

**Results:** We identified seven monocyte clusters. Their abundances varied between cirrhosis stages, confirming previously reported changes i.e. reduction in CD14^low^CD16^++^ and emergence of M-MDSC in advanced stages. DE genes between health and disease and among stages were detected, including for the first time CD52. CD52-expression on monocytes significantly enhanced throughout compensated and NAD cirrhosis. Heretofore the biological significance of CD52-expression on monocytes remained unknown. CD52^high^CD14^+^CD16^high^HLA-DR^high^ monocytes in patients with cirrhosis revealed a functional phenotype of active phagocytes with enhanced migratory potential, increased cytokine production, but poor T cell activation. Following acute decompensation (AD), CD52 was cleaved by elevated phospholipase C (PLC), and soluble CD52 (sCD52) was detected in the circulation. Inhibition and cleavage of CD52 significantly suppressed monocyte functions *ex vivo* and *in vitro*, and the predominance of immunosuppressive CD52^low^ circulating monocytes in patients with AD was associated with infection and low transplant-free survival.

**Conclusion:** CD52 may represent a biologically relevant target for future immunotherapy. Stabilising CD52 may enhance monocyte functions and infection control in the context of cirrhosis, guided by sCD52/PLC as biomarkers indicating immuneparesis.

**Lay summary:** Recurrent infections are a major cause of death in patients with liver cirrhosis. A fundamental understanding of the mechanisms that suppress immune responses in patients with cirrhosis is lacking, but required for the development of strategies to restore innate immunity in cirrhosis patients and prevent infection. The current study identified a novel marker for deficient immune responses and a potential target for such a future immune-based therapy.

Graphical Abstract
scRNA-seq identified seven circulating monocyte states, changing in cirrhosis patients at different stages of disease. Circulating monocytes overexpress CD52 in cirrhosis, but are absent in AD/ACLF due to PLC. CD52-expressing monocytes show high capability for phagocytosis, cytokine production, adhesion and migration potential and T cell suppression. Created with BioRender.com

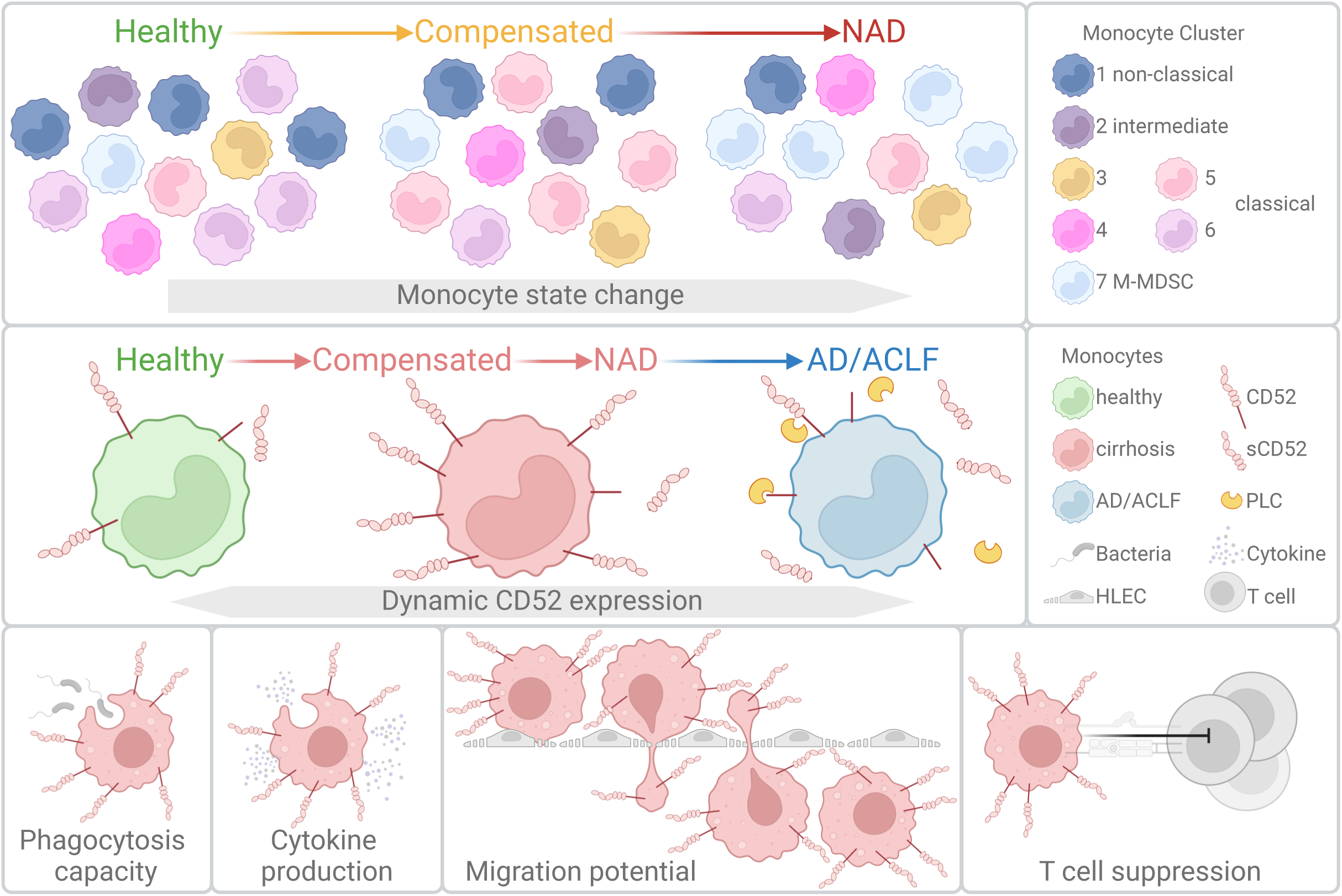

## Introduction

Cirrhosis of the liver is a systemic inflammatory condition initiated by chronic liver injury and scarring, destruction of the distinct liver vasculature, loss of hepatic function and consequently significant systemic metabolic and immunological changes [1, 2]. Onset and recurrence of bacterial infections in patients suffering from cirrhosis relate to the significant morbidity and mortality being the most common “precipitating factors” eliciting decompensation, hospitalisation and significantly increase the risk of death [3]. The sequence of pathophysiological events evolving over years remains incompletely understood. Particularly, the understanding of the onset and evolution of systemic immuneparesis, in relation to the different stages of disease, i.e. compensated and decompensated cirrhosis, acute decompensation (AD) and liver failure (ACLF) may enable future immunotherapy to prevent deterioration and bridge to regeneration [4] or transplantation.

Immuneparesis in cirrhosis involves an attenuated function of diverse innate and adaptive immune cells, and soluble factors in multiple compartments [5]. Along with disease progression, circulating monocytes from patients with cirrhosis lose their regular differentiation and function. In AD/ACLF stages immunosuppressive states i.e. low HLA-DR expressing monocytic myeloid derived suppressor cells (M-MDSC) and MERTK-expressing monocytes prevail [6, 7]. Yet, M-MDSC already occur in earlier compensated stages as well as immune-modulatory AXL-expressing monocytes [5, 8]. We hypothesise that unrecognised monocyte states may modulate immune responses in cirrhosis, and therefore, pursued a systematic unbiased approach to identify states of monocyte differentiation in relation to disease stages.

CD52 is a small glycosylphosphatidylinositol-anchored protein, expressed on all leukocytes. CD52 has mostly been described on T cells, limiting T cell activation [9]. A monoclonal antibody against CD52 (alemtuzumab) is in clinical use for the treatment of haematologic malignancies and multiple sclerosis [10]. It depletes lymphocytes by an antibody-dependent, cell-mediated cytotoxicity [10], aiming at beneficial reconstruction of the immune system. The function of CD52-expression on monocytes has barely been described. While soluble CD52 (sCD52) suppressed inflammatory cytokine responses by monocytes, CD52 deletion exacerbated inflammation [11]. Up-regulation of CD52 and co-localisation with its receptor SIGLEC-10 has been shown on monocytes and quiescent stem-like cells in the context of Acute myeloid leukemia (AML) [12]. Of note, CD52-expression on lymphocytes is a prognostic biomarker for sepsis and correlated with improved survival [13].

In this work, hypothesising, that heretofore unrecognised monocyte states may modulate immune responses in cirrhosis, we pursued a systematic unbiased approach to identify states of monocyte differentiation in relation to disease stages.

## Material and Methods

### Patients and Sampling

A cohort of 90 patients was identified at the University Hospital Basel and the Cantonal Hospital St. Gallen, Switzerland from June 2019 until June 2023. Patients with biopsy-proven cirrhosis were recruited during outpatient consultation or hospital admission for decompensation (Table S1) or liver resection surgery (Table S2), respectively. Cirrhosis patients were categorised according to D’Amico [14] into compensated [n=31], not-acute-decompensated (NAD) [n=17] and AD/ACLF [n=15]. We also included healthy controls (HC [n=27]). Resection tissue samples were categorised into cirrhosis (n=12) and morphologically normal controls (n=16). Exclusion criteria were age younger than 18 years and evidence of HIV infection. Blood samples were obtained for *ex vivo* analysis of monocyte differentiation and function, plasma/serum, and peripheral blood mononuclear cells (PBMC) were stored. Liver tissue specimens were obtained from liver resection surgery for *ex vivo* analysis of liver macrophage differentiation and function. Patients were followed up for 1 year for adverse events. The study was approved by the local ethics committees (EKSG 15/074/EKNZ 2015-308, BASEC-ID 2019-00816, BASEC-ID 2019-02118) and recorded in the clinical trial register ClinicalTrials.gov (NCT04116242) and Swiss National Clinical Trials Portal (SNCTP000003482).

### Clinical, Haematologic, and Biochemical Parameters

Full blood count, liver and renal function tests, and clinical variables were obtained by the clinician and entered into a database.

### Cell isolation

Monocytes, T cells and human hepatic macrophages were isolated as previously described [8, 15].

### Single cell RNA sequencing methods

Blood samples from a subset of 16 patients, including 6 healthy donors, 5 patients with compensated cirrhosis (one later excluded, see below) and 5 with decompensated cirrhosis, were processed for scRNA-seq, across 8 batches from July 2019 to August 2021. Enriched monocytes isolated by magnetic cell separation (MACS) were stained for viability and monocyte markers (CD14, CD16, HLA-DR) and sorted using BD FACSMelody Cell Sorter. For each sample, an estimated count of 20,000 monocytes were loaded onto a single 10X channel, for a targeted recovery of 10,000 cells. cDNA and library preparation was performed with a Single Cell 3’ v3 Reagent Kit (10X Genomics) according to the manufacturer’s instructions. Sequencing was performed on an Illumina platform to produce 90nt-long R2 reads. The dataset was analyzed by the Bioinformatics Core Facility, Department of Biomedicine, University of Basel (Supplementary Methods).

### Flow Cytometry based Phenotyping

Phenotyping of peripheral blood monocytes, neutrophils, PBMC and hepatic tissue macrophages was undertaken using flow cytometry (Table S3) (LSR Fortessa Cell Analyzer, BD Biosciences) as previously described [6]. Monocytes from HC expressing low CD52 levels (Median: 32.4%, 348.5MFI) (<40% / <600MFI CD52-expressing monocytes) and monocytes from patients with cirrhosis expressing high CD52 levels (Median: 75.80% / 72.65%, 644MFI / 496MFI) (>60% / >600MFI CD52-expressing monocytes) were identified and further processed for phagocytosis capacity, mixed lymphocyte reaction, adhesion and migration assay, *in vitro* inhibition with alemtuzumab and plasma conditioning.

### *In vitro* assessment of monocyte function

Phagocytosis capacity, cytokine responses of monocytes upon Toll-like receptor (TLR) stimulation and T cell proliferation activation was undertaken using flow cytometry as previously described [7, 8]. Evaluation of monocyte adhesion to ICAM-1, ICAM-2, VCAM-1, and CD52 inhibition was undertaken as previously described [16]. For plasma conditioning CD52^low^ monocyte samples and CD52^high^ monocyte samples were incubated with plasma as previously described [7]. Effect of phospholipase C (PLC) on CD52-expression on monocytes was assessed as previously described [17].

### Migration Assay in Static Conditions through Human Liver-derived Endothelial Cells

Transwell inserts were coated with collagen (30%) and 1×10^6^ human liver-derived endothelial cells (HLEC) were plated on the inserts. Once confluency was reached 2×10^6^ monocytes were plated and migrated monocytes were assessed after 24h by flow cytometry [18].

### Generation of THP-1 Cell Line Stably Expressing CD52

pLV[Exp]-CMV>hCD52[NM_001803.3] vector was designed and ordered from VectorBuilder. Packaging plasmids psPAX2 (from Didier Trono, Addgene plasmid #12260) and pMD2.G (from Didier Trono, Addgene plasmid #12259) and were used for production of lentivirus [19]. After sorting (BD FACSMelody) around 60% of THP-1-hCD52 cells expressed hCD52, while CD52-expression level of non-transduced counterparts was 0%.

### ELISA assays

sCD52 and PLC were measured using enzyme-linked immunosorbent assay (ELISA) in plasma as previously described [16] and according to the manufacturers protocol (antibodies.com, Human PLCG 2 Kit Elisa, Cat.: A75730).

### Statistical Analysis

Statistical evaluation and graph design was performed in Prism (GraphPad, La Jolla, CA; version 9.5.1). Data are expressed as the median/interquartile range unless otherwise specified. For data that did not follow a normal distribution, the significance of differences was tested using the Mann-Whitney-, Wilcoxon-, Kruskal-Wallis- or Friedman-test; Spearman correlation coefficients were calculated and p<0.05 values were considered statistically significant. Data are presented as Column or box plots showing median with 10-90 percentile and all points. *p<0.05/**p<0.01/***p<0.001/****p<0.0001

## Results

### scRNA-seq revealed distinct abundance of monocyte clusters in compensated and decompensated cirrhosis

Our scRNA-seq dataset gathered monocytes from 15 healthy and cirrhosis individuals (Fig. 1A; Fig. S1A-C). After quality filtering (Fig. S1D) and correction for patient-specific effects (Fig. S1C), a total of 100,162 monocytes were partitioned into 7 clusters visualized onto a UMAP embedding (Fig. 1B), which were annotated using unbiased comparisons to references atlases (Fig. S1D), and the expression of cluster specific genes (Fig. 1C).

**Figure 1:**
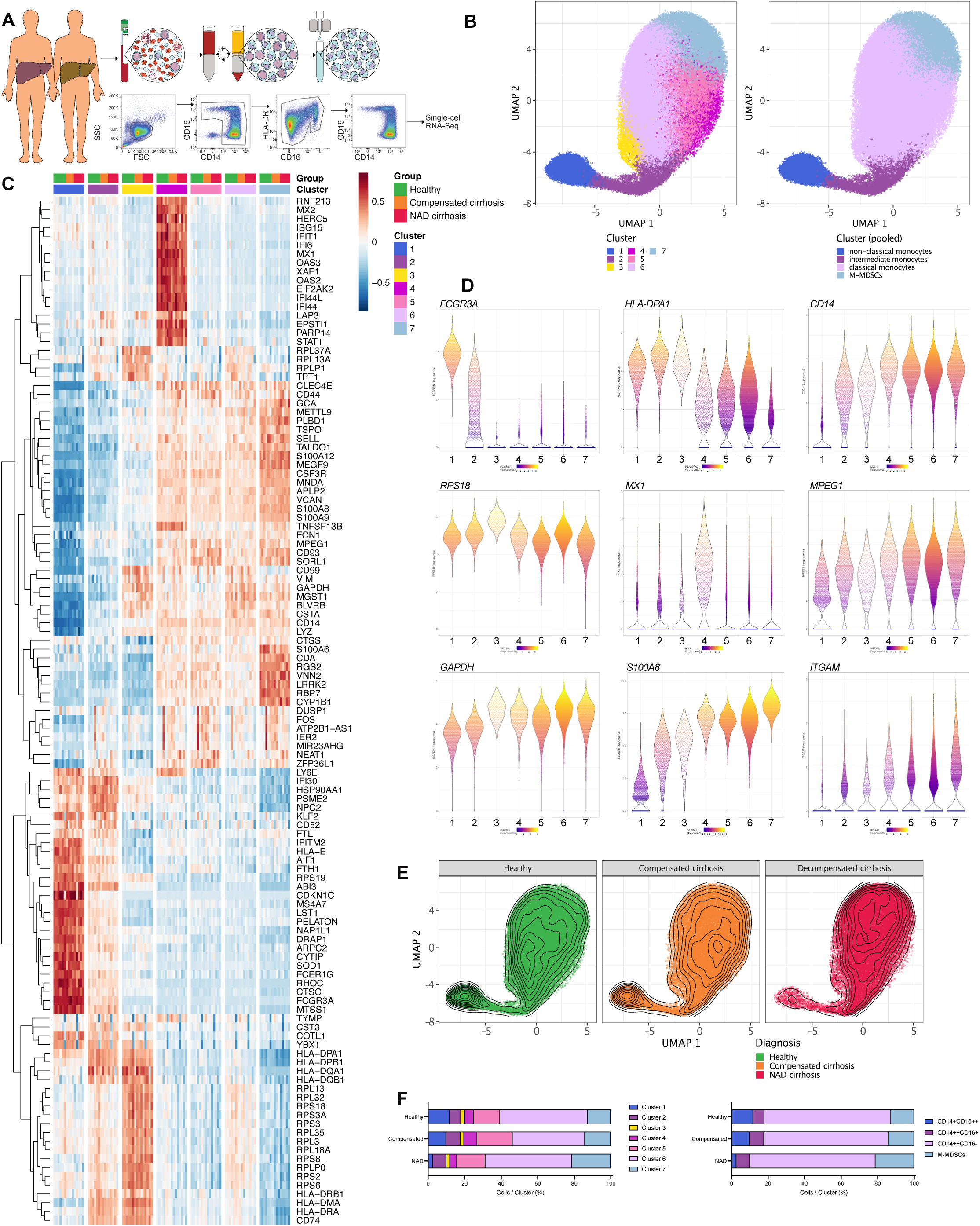
Single-cell atlas of human circulating monocytes. **A:** Overview of monocyte isolation from HC (n=6), compensated (n=4) and NAD (n=5) cirrhosis patients. **B:** Heatmap of cluster-defining marker genes. The log-normalized expression levels were averaged across cells from the same cluster and sample, and averaged values were then centred and scaled per gene. **C:** Final clustering of monocytes visualized on a UMAP embedding. **D:** Gene expression of selected cluster-defining markers across cells. **E:** Density of monocytes across disease state. **F:** Cluster frequencies across disease state.

Cluster 1 was defined by Fc-gamma receptors (e.g., *FCGR3A, FCER1G*), and represented non-classical monocytes (CD14^+^CD16^++^). Expression of MHC class II members (e.g., *HLA-DRB1, HLA-DRA*) was enriched in cluster 2 and represented intermediate monocytes (CD14^++^CD16^+^). Clusters 3 to 6 represented classical monocytes (CD14^++^CD16^−^). In cluster 3 MHC class II and ribosomal protein genes (e.g., *RPL13, RPS18)* were highly expressed. Interferon induced proteins (e.g., *MX1, IFIT1)* defined cluster 4. Cluster 5 and 6 showed similar expression patterns, although cluster 5 expressed genes involved in phagocytic processes (e.g., *MPEG1, CD93*). Cluster 6 displayed slightly elevated expression of ribosomal proteins (e.g., *RPS3A, RPL3, RPL37A*) and genes related to cell degradation and recycling (e.g., *CD99, VIM, GAPDH, MGST1I*). Finally, cluster 7 was enriched in S100 protein genes (e.g., *S100A8, S100A9*), *CD84, ITGAM, CD33* but low in MHC II, and thus likely corresponds to M-MDSC (Fig. 1C, D).

Overall, we thus observed functionally distinct monocytic subsets, which notably included the previously described separation between CD16^high^ and MHC II^high^ expressing monocytes on one side and CD14high monocytes on the other side.

Monocytic clusters relative frequencies were then compared (Fig. 1E, F), but a large variability was present across patients within each group (Fig. S1). This was likely the reason for the absence of significant differences between compensated cirrhosis and healthy subgroups. However, the direct comparison of NAD cirrhosis to healthy showed a significant decrease in frequency of cells from cluster 1 (non-classical) and a significant increase in frequency of cells from cluster 7 (M-MDSC), as previously described [7, 8] (Fig. 1E, F).

### Transcriptome changes of monocytes in cirrhosis

Differentially expression analysis between patient subgroups was stratified by cluster to eliminate the potential effects of the above-described cluster frequency differences. Notably differentially expressed (DE) genes were identified in non-classical (cluster 1), intermediate (cluster 2), classical monocytes (cluster 3-6) and M-MDSC (cluster 7) (Fig. 2A / S2).

**Figure 2:**
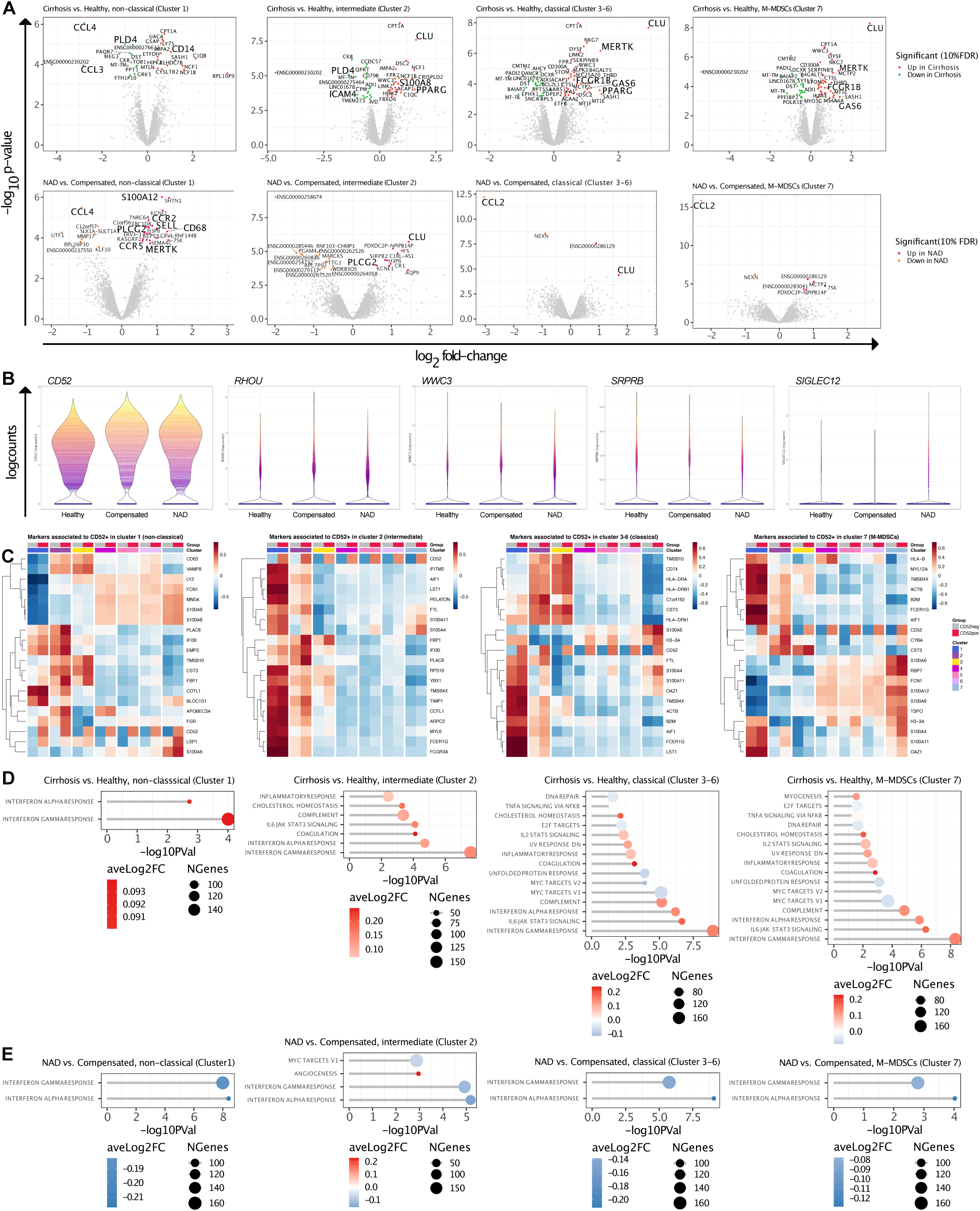
Identification of differential Gene expression in monocytes in liver cirrhosis. **A:** Volcano plots showing DE genes in cirrhosis (n=9) vs. HC (n=6) (top row) and NAD (n=5) vs. compensated (n=4) cirrhosis (bottom row) in different clusters. **B:** Violin plots of DE gene expression. **C:** Heatmap of DE genes comparing *CD52^high^* vs. *CD52^low^* monocytes. GSEA of **D:** cirrhosis vs. HC and **E:** NAD vs. compensated.

Comparing cirrhosis to HC samples in cluster 1 we found that pro-inflammatory cytokines (*CCL3, CCL4*) were downregulated, and genes involved in apoptosis (*UACA*), complement system (*C1QB*), lipopolysaccharide (LPS) sensing (*NCF1*), TLR4 signalling (*SASH1*) and antigen processing (*LY75*) were upregulated. In cluster 2 GTPase activator (*ACAP1*) and calprotectin (*S100A8*) were upregulated in cirrhosis. In CD14^high^ clusters (3-6) *DANCR* and *PADI2* were decreased, *NKG7, CD300A, GAS6* and *LIMK2* upregulated in cirrhosis. Noteworthy, *NKG7* is expressed by monocytes differentiating into macrophages [20] and *CD300A* blocks TLR4-MyD88 induced signalling [21]. *GAS6* is a TAM receptor ligand to which family MERTK and AXL belong to [22, 23]. Also, the TAM receptor *MERTK* was upregulated in cirrhosis as described before [7] (Fig. 2A). Most of the DE genes detected in the cluster 7 (M-MDSC): *CD300A, CLU, CMTM2, CPT1A, DYSF, IKBKE, NKG7* were also detected in other clusters, but *THBD*, binding thrombin and activating protein C was upregulated only in this cluster (Fig. 2A, upper panel). Secondly, we analysed genes differentially regulated in NAD compared to compensated cirrhosis. In NAD in most clusters *CCL2* (chemokine) and *NEXN* (pyroptosis inhibitor) were downregulated, and *CLU* and *CPVL* (STAT1 inhibitor) upregulated. In cluster 1 *S100A12, CCR2, SELL, CD68* and *PLCG2* were upregulated in NAD indicating inflammation and migration. Of note, the receptor *CCR2* was upregulated in NAD, *CCL2* and *CCL4* downregulated. In cluster 2 coagulation factor *F5* was upregulated, while the PKC substrate *MARCKS* and ribosomal protein *RPL7P9* were downregulated in NAD (Fig. 2A lower panel).

The analyses suggested that genes of the phosphatidylinositol signalling pathway, i.e., *PLCG2, PLD4, IMPA2,* were among differentially expressed in monocytes of cirrhosis patients compared to HC (Fig. 2A). Interestingly also *CD52,* belonging to the family of glycosylphosphatidylinositol (GPI)-anchored proteins, was found consistently more expressed in cirrhosis than in HC (although this difference was never significant, e.g., cirrhosis vs. HC in cluster 1: log_2_FC=0.59, p-value=0.053, FDR=100%). Other GPI-anchored proteins like CD24 were upregulated upon WNT signalling [24]. DE gene analysis found genes involved in WNT signalling pathway (*WWC3, RHOU, SRPRB*) deregulated in cirrhosis. Furthermore, CD52 is shed by PLC in T cells [25], and its gamma isoform (*PLCG2*) was detected upregulated in NAD. Moreover, sCD52 binds to SIGLEC10 to suppress T cell activation, and DE gene analysis identified *SIGLEC12* (Fig. 2B). Thus, we identified *CD52* as a gene with putative central role in the regulation of monocyte function in cirrhosis affecting genes involved in WNT and SIGLEC signalling pathways.

To gain first insights into *CD52*-expression in monocytes, *CD52^pos^* monocytes were compared to *CD52^neg^*, which revealed that expression of *S100* proteins was associated with *CD52^pos^* monocytes in all clusters. *CD52^pos^*monocytes showed higher expression of *IFI30* and *MNDA* in cluster 1 (non-classical), interferon-induced proteins (e.g., *IFITM2, IFI30*) and Fc-receptors (e.g., *FCER1G, FCGR3A*) in cluster 2 (intermediate), MCH II members (e.g., *CD74, HLA-DRA, HLA-DRB1*) in classical monocytes and *S100* proteins, but also *FCER1G* and *HLA-B* in cluster 7 (M-MDSC) (Fig. 2C).

For the identification of common up- or downregulated pathways, gene set enrichment analysis (GSEA) was performed using the Hallmark Signatures gene sets.

In cirrhosis monocytes inflammatory responses like interferon-α/-γ, IL-2/IL-6 signalling, phagocytosis and the complement system were upregulated. Interestingly, also pathways involved in coagulation, cholesterol- and lipid-homeostasis were upregulated in cirrhosis, while DNA repair and MYC targets were downregulated compared to HC (Fig. 2D). Comparing pathways differentially regulated in NAD compared to compensated cirrhosis, however, inflammatory responses like interferon-α/-γ and TNFα signalling were downregulated (Fig. 2E).

Using Gene Ontology Term (data not shown) data set for GSEA multiple pathways regulating T cell proliferation/differentiation and cellular adhesion were upregulated, which were previously found to be affected by the overexpression of *CD52* [25].

### Upregulation of CD52 on monocytes indicates lack of infection and survival

We analysed peripheral blood monocytes from cirrhosis patients and HC to confirm *CD52* upregulation at the protein level. Monocyte count was unchanged in cirrhosis (Fig. 3A). CD52^high^ expressing monocytes were expanded in patients with compensated and NAD cirrhosis, compared to HC, but not AD/ACLF (Fig. 3B). Flow cytometry analysis of neutrophils revealed a significant downregulation of CD52 in AD/ACLF compared to compensated/NAD stages, but not other PBMC (Fig. 3C).

**Figure 3:**
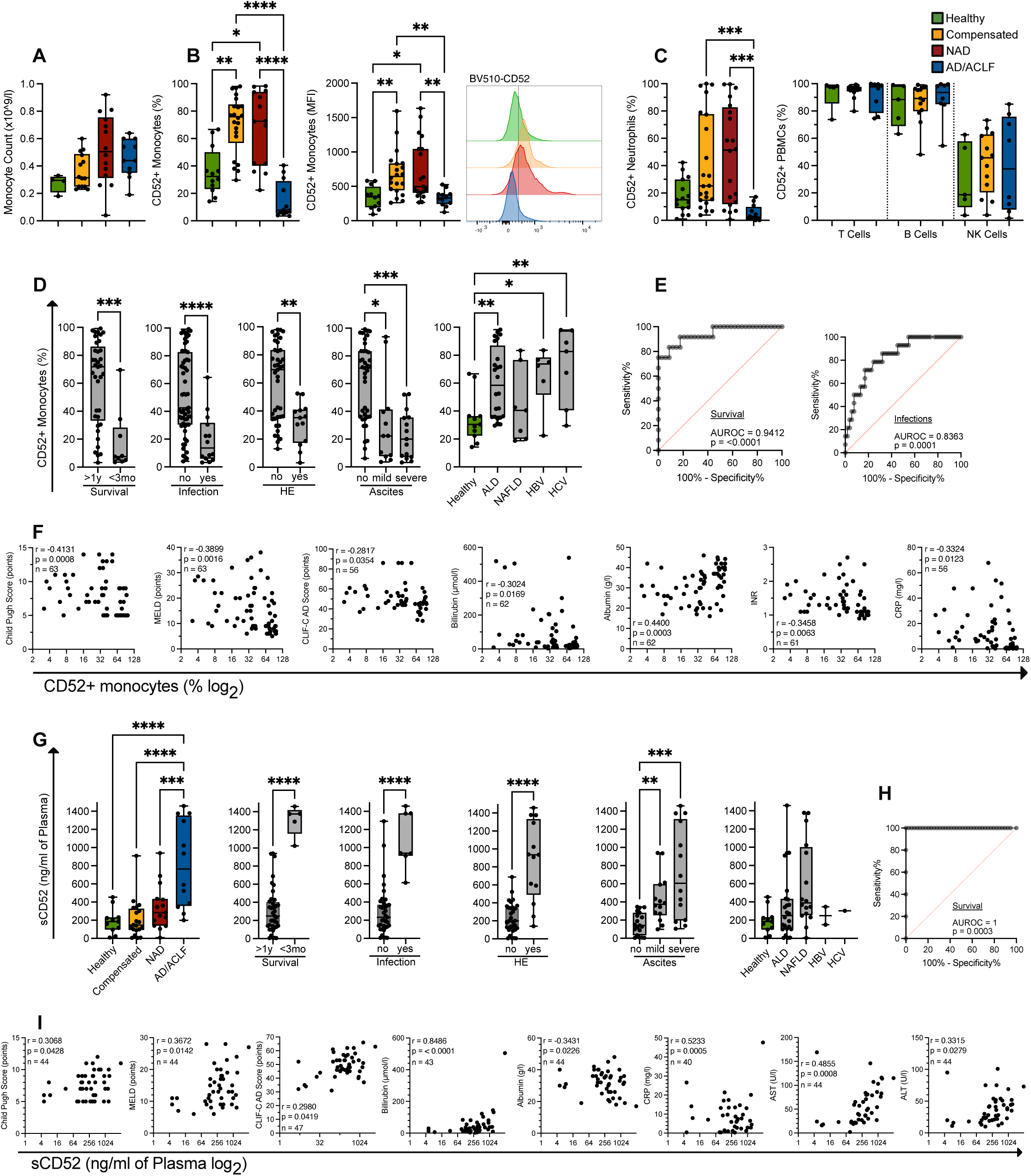
Clinical relevance of CD52 expression on monocytes. **A:** Monocyte counts. CD52-expression on **B:** circulating monocytes (% and MFI) with representative FACS-histograms, **C:** Neutrophils (%), T-cells, B-cells and NK cells (%) at different cirrhosis stages. CD52-expression on monocytes **D:** related to survival, infection, HE, ascites and cirrhosis aetiology, **E:** predicts 1-year mortality, infection and **F:** correlates negatively with Child Pugh, MELD, CLIF-C-AD, bilirubin, albumin, INR, CRP. Plasma levels of sCD52 **G:** at different cirrhosis stages, related to survival, infection, HE ascites, cirrhosis aetiology, **H:** predicts survival and **I:** correlate with Child Pugh, MELD, CLIF-C-AD, bilirubin, albumin, CRP, AST, ALT. *p<0.05/**p<0.01/***p<0.001/****p<0.0001 (Kruskal-Wallis-, Friedman-, Mann-Whitney-test, Spearman correlation coefficient)

Regarding its clinical significance, CD52-expression on monocytes from cirrhosis patients was independent of the underlying aetiology and associated with transplant free survival (TFS), development of infectious complications, hepatic encephalopathy (HE) and ascites (Fig. 3D); it predicted 1-year TFS (AUROC 0.9412) and the development of infection (AUROC 0.8363) (Fig. 3E). Given CD52^high^ monocytes occurred in compensated and NAD cirrhosis, but not AD/ACLF (Fig. 3B), the proportion of CD52-expressing monocytes negatively correlated with disease severity scores (Child Pugh, MELD, CLIF-AD) and individual parameters of liver function (bilirubin, Albumin, INR, CRP) (Fig. 3F).

To determine whether CD52 might be shed from monocytes when patients progress to AD/ACLF stage of cirrhosis, sCD52 in plasma was determined, and indeed sCD52 plasma levels were elevated significantly only in AD/ACLF (Fig. 3G). High sCD52 plasma levels were associated with low TFS, infection, HE and ascites, independent of the underlying aetiology (Fig. 3G). Plasma levels of sCD52 predicted 1-year TFS (AUROC 1, Fig. 3H). plasma sCD52 positively correlated with disease severity scores (Child Pugh, MELD, CLIF-AD) and parameters of liver function (Fig. 3I).

### CD52^high^ monocytes implicate macrophage-like activated phagocytes dampening adaptive immune stimulation

CD52-expression levels were assigned to distinct monocytic subsets (Fig. 4A). Classical CD14^high^ monocytes increased in cirrhosis, while non-classical CD16^high^ decreased significantly in AD/ACLF compared to HC (Fig. 4B). In classical monocytes CD52-expression increased in compensated and NAD cirrhosis, but decreased significantly in AD/ACLF. In intermediate and non-classical monocytes, the CD52-expression decreased in AD/ACLF (Fig. 4C).

**Figure 4:**
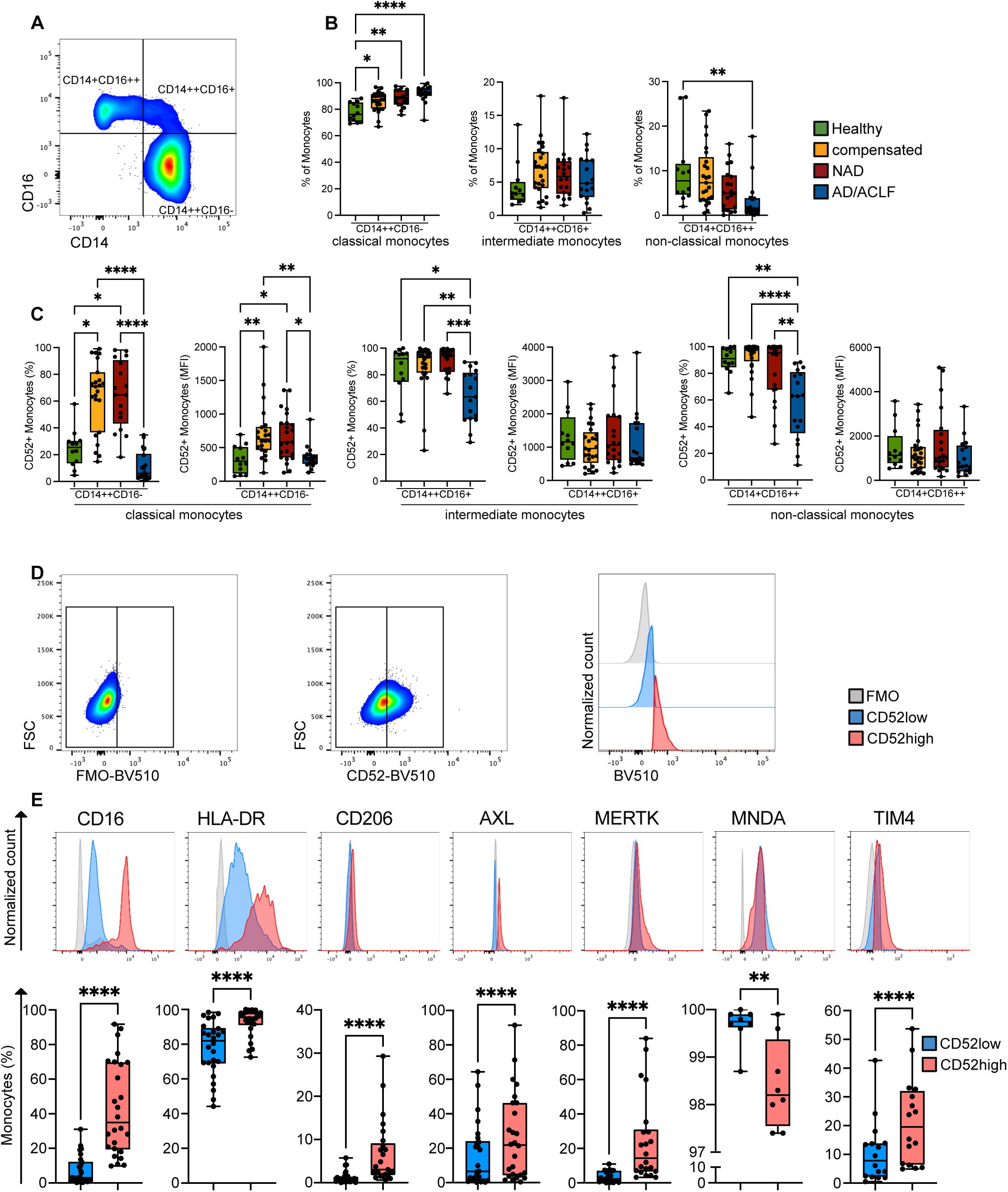
Phenotypic characterisation of CD52high monocytes. **A:** Gating strategy for monocytic subset identification (classical=CD14^++^CD16^−^, intermediate=CD14^++^CD16^+^, non-classical=CD14^+^CD16^++^). **B:** Monocyte count across cirrhosis stages. **C:** CD52-expression (% and MFI) of monocytic subsets across cirrhosis stages. **D:** Gating strategy and representative histograms to identify CD52^high^ and CD52^low^ monocytes. Forward scatter (FSC), fluorescence minus one (FMO) **E:** Immunophenotyping of CD52^high^ and CD52^low^ monocytes in cirrhosis. *p<0.05/**p<0.01/***p<0.001/****p<0.0001 (Kruskal-Wallis-, Wilcoxon tests)

CD52^high^ monocytes (Fig. 4D) were CD14^+^CD16^high^HLA-DR^high^ indicating a mature monocyte population and showed higher levels of macrophage markers CD206 and TIM-4 [4, 26] and TAM receptors AXL and MERTK [6, 8], but lower levels of the circulation marker MNDA [27] (Fig. 4E).

Given the biological function of CD52 on monocytes is not well known, we detailed functional characteristics of circulating CD52^high^ and CD52^low^ monocytes from patients with cirrhosis. Phagocytosis was similar among patients with cirrhosis and HC when challenged with *E.coli* particles, but increased towards *S.aureus* in monocytes from cirrhosis patients *ex vivo* (Fig. 5A). CD52^high^ monocytes exhibited enhanced phagocytosis capacity compared to their CD52^low^ counterparts (Fig. 5B). Baseline production of TNF-α and IL-10 were significantly increased in monocytes from cirrhosis patients compared to HC, IL-6 levels were similar. Stimulation with TLR ligands LPS and synthetic triacylated lipopeptide Pam3CSK4 (Pam3) revealed lower TNF-α/IL-6 production in monocytes from cirrhosis patients compared to HC, but increased IL-10 production upon Pam3 (Fig. 5C). Noteworthy, CD52^high^ monocytes showed higher TNF-α, IL-6, IL-10 production compared to CD52^low^ (Fig. 5D).

**Figure 5:**
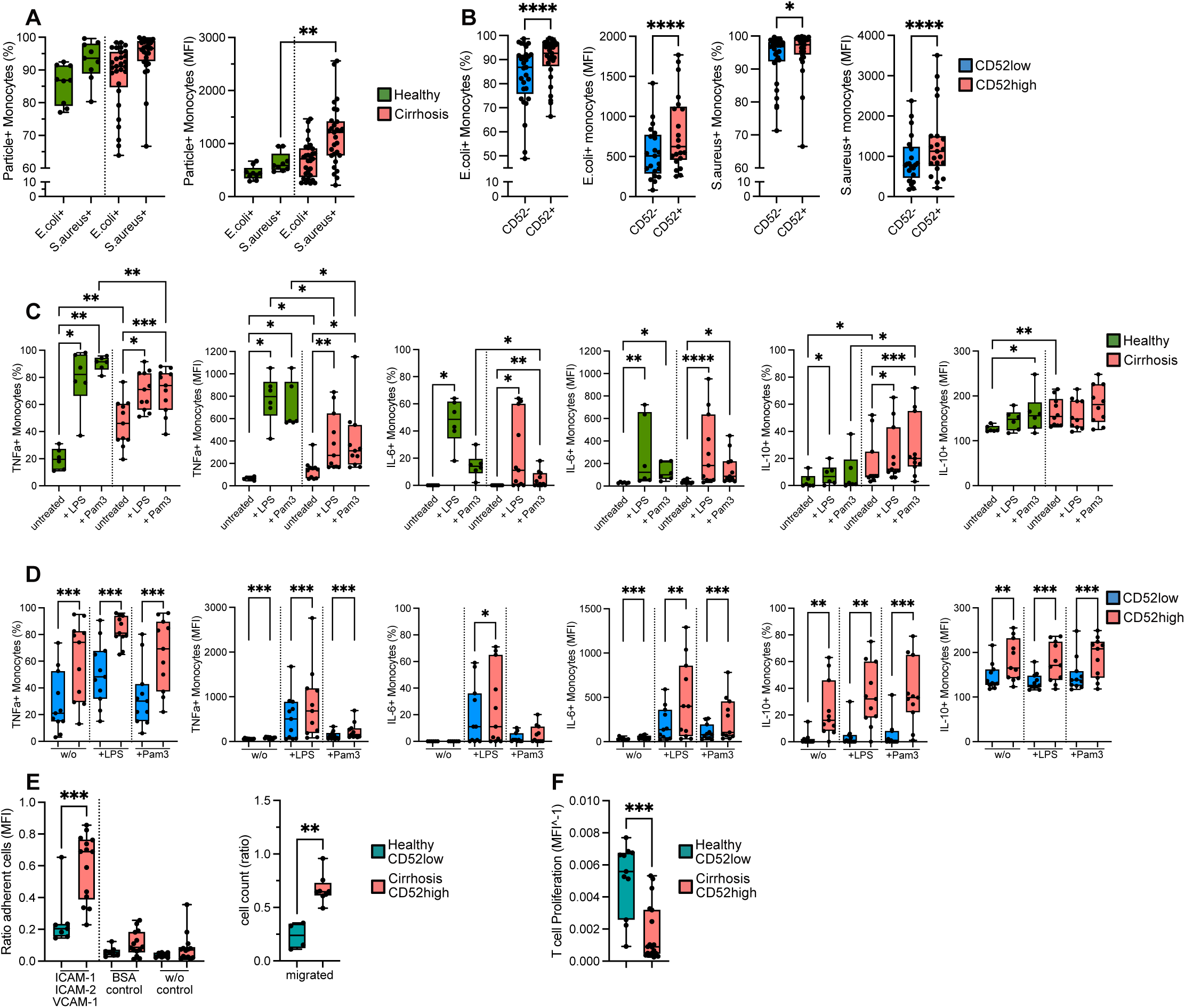
Functional characteristic of CD52high monocytes *ex vivo*. Phagocytosis capacity (% and MFI) of monocyte in **A:** HC vs. cirrhosis and **B:** CD52^high^ vs. CD52^low^ from cirrhosis. Cytokine production (% and MFI) upon TLR stimulation of monocytes in **C**: HC vs. cirrhosis and **D:** CD52^high^ vs. CD52^low^ from cirrhosis. **E:** Adhesion, migration behaviour (MFI and ratio) and **F:** T-cell proliferation (MFI^-1) in co-culture with monocytes, of previously defined CD52^high^ vs. CD52^low^ samples. *p<0.05/**p<0.01/***p<0.001/****p<0.0001 (Kruskal-Wallis-, Wilcoxon-, Friedman-, Mann-Whitney-test)

In inflammatory conditions such as liver injury circulating blood monocytes infiltrate tissues e.g. the liver and give rise to monocyte derived macrophages [28]. We observed enhanced adhesion and migration capabilities of monocytes from cirrhosis patients (Fig. 5E).

Furthermore, we investigated the potential of monocytes to modulate T cell proliferation [8] and observed that proliferation was dampened in presence of CD52^high^ cirrhotic monocytes compared to CD52^low^ healthy monocytes *ex vivo* (Fig. 5F).

### Modulation of CD52 expression *ex vivo* induces significant functional changes

Given the biological significance of CD52 upregulation on monocytes is unknown we developed a THP-1 cell line overexpressing human CD52 (hCD52; THP-1-hCD52) (Fig. 6A). Lentiviral transduction of THP-1 cells with hCD52 resulted in a secondary antigen with an expression pattern over a continuum (Fig. 6B). Similar to CD52^high^ monocytes from cirrhosis patients, THP-1-hCD52 cells showed increased expression of activation markers CD16, HLA-DR, CD206, TIM-4, AXL, MERTK compared to THP-1; relating to the putative biological relevance, the expression level of CD16, CD206, TIM-4, AXL were in general below 5% (Fig. 6C). Similar to patient derived monocytes, baseline and TLR-stimulated cytokine production was increased in THP-1-hCD52 cells, while not significant for IL-6 production (Fig. 6D).

**Figure 6:**
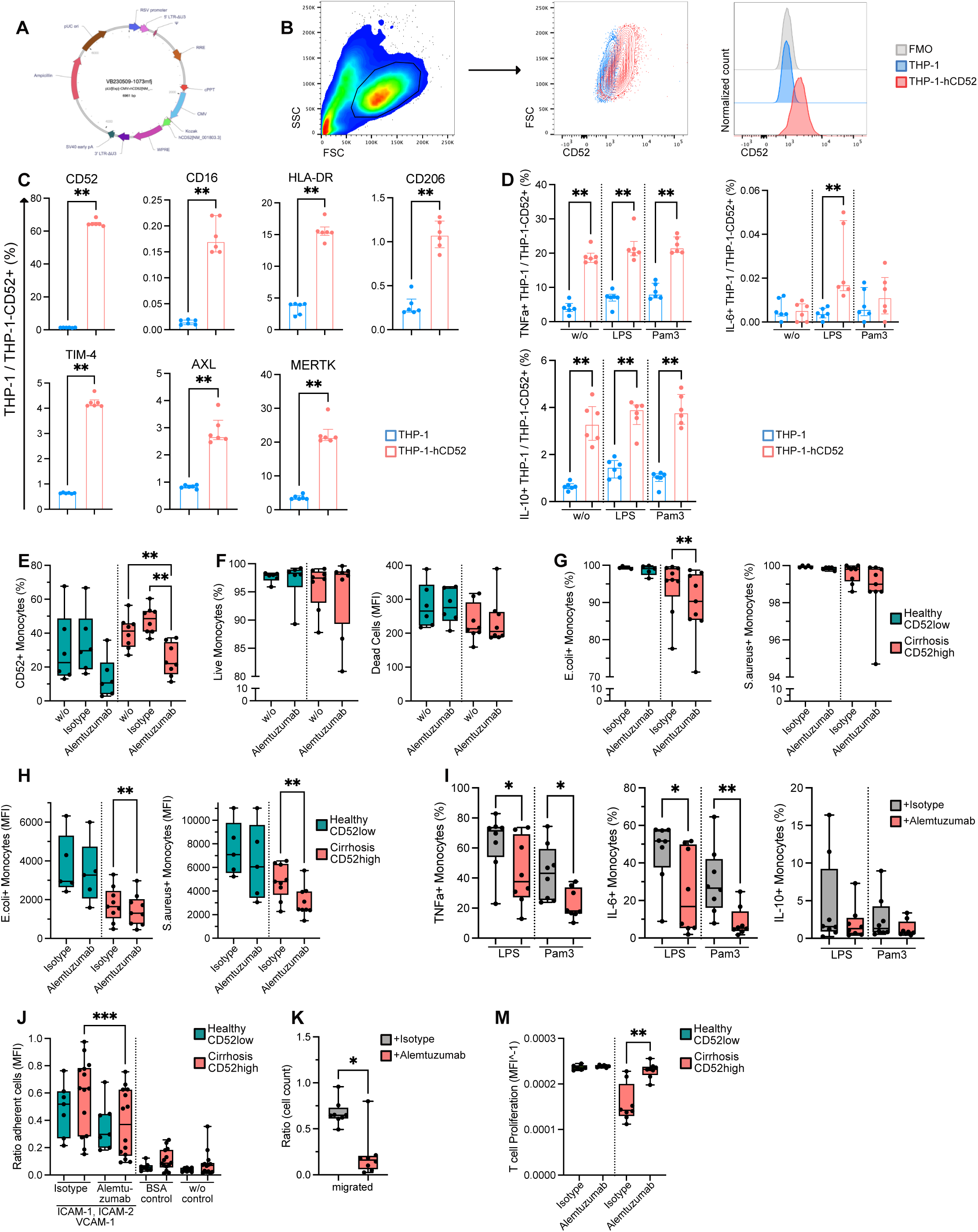
CD52 overexpression in THP-1 cells and inhibition in monocytes *in vitro*. **A:** Plasmid used for lentiviral transduction. **B:** Representative FACS plot and histograms to identify THP-1-hCD52 cells. Forward scatter (FSC), side scatter (SSC), fluorescence minus one (FMO). **C:** Immunophenotyping of THP-1-hCD52 and THP-1 cells. **D:** Cytokine production upon TLR stimulation of THP-1-hCD52 and THP-1 cells. Effect of CD52 inhibition by alemtuzumab treatment on monocytes of previously defined CD52^high^ vs. CD52^low^ samples: **E:** CD52-expression (%), **F:** survival (% and MFI), **G:, H:** phagocytosis capacity (% and MFI), **I:** cytokine production (%) upon TLR stimulation, **J:** adhesion and **K:** migration potential (ratio). **M:** T-cell proliferation (MFI^-1) in co-culture with monocytes from previously defined CD52^high^ patients and CD52^low^ HC samples. *p<0.05/**p<0.01/***p<0.001/****p<0.0001 (Mann-Whitney-, Friedman-, Wilcoxon-, Kruskal-Wallis test)

Furthermore, the effects of CD52 inhibition were examined. Treatment of isolated monocytes with alemtuzumab, reduced CD52 surface expression on monocytes from patients (CD52^high^), compared to isotype control and untreated cells, but not monocytes from HC (CD52^low^) (Fig. 6E). Viability of alemtuzumab treated monocytes did not change (Fig. 6F).

Functionally, alemtuzumab treatment lead to reduced phagocytosis of *E.coli* and *S.aureus* by monocytes from cirrhotic patients but not HC (Fig. 6G, H). Cytokine production of TNF-α and IL-6 upon TLR stimulation (LPS, Pam3) of circulating CD52^high^ monocytes was reduced upon alemtuzumab treatment (Fig. 6I). Blocking of CD52 on monocytes also decreased adhesion and migration capabilities of CD52^high^ monocytes from cirrhosis patients when compared to controls (Fig. 6J, K). The suppressive phenotype of cirrhotic CD52^high^ monocytes towards T cell activation was inverted almost to a level similar to CD52^low^ monocytes from HC (Fig. 6M).

### Plasma components regulate CD52 expression and phagocytosis capacity

We further evaluated possible mechanisms modulating CD52-expression on monocytes. Pathophysiologically, bacterial translocation facilitates microbial products accessing the circulation in cirrhosis patients [28], and may cause changes in the differentiation of circulating monocytes. Conditioning of healthy donor monocytes with plasma samples from patients at different cirrhosis stages revealed that CD52 was upregulated in plasma from compensated and NAD cirrhosis, but not AD/ACLF patients (Fig. 7A). Co-culture of CD52^high^ monocytes from cirrhosis patients showed downregulation of CD52-expression after treatment with AD/ACLF patients’ or HC plasma (Fig. 7B). To investigate if bacterial products were the plasma component causing CD52 upregulation in compensated and NAD cirrhosis, the percentage of CD52-expressing monocytes was assessed after stimulation with bacterial particles and TLR ligands (LPS, Pam3). CD52-expression was significantly increased on monocytes after phagocytosis of *E.coli* and *S.aureus* bioparticles (Fig. 7C), but treatment of monocytes with LPS alone was insufficient to upregulate CD52-expression (Fig. 7D). Noteworthy, also migrated monocytes from cirrhosis patients showed higher expression of CD52 compared to non-migrated (Fig. 7E). Furthermore, AD/ACLF patients’ plasma dampened CD52-expression and also phagocytosis capacity of monocytes from HC and cirrhosis (Fig. 7F, G).

**Figure 7:**
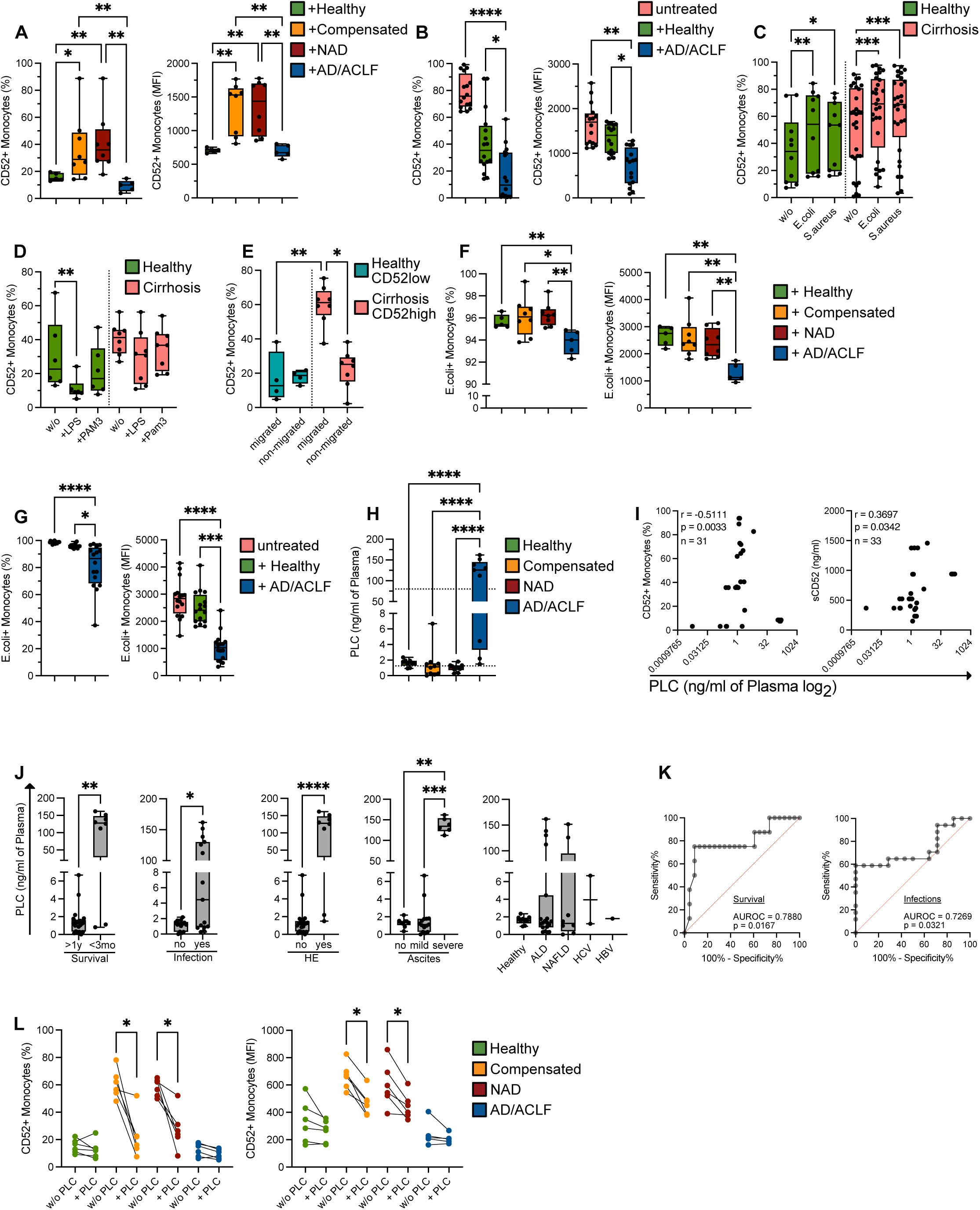
Effect of plasma conditioning on CD52high monocytes *in vitro*. CD52 expression (% and MFI) of **A:** HC monocytes and **B:** previously defined CD52^high^ monocytes from cirrhosis patients upon plasma conditioning from different disease stages. CD52 expression (%) on monocytes of HC and cirrhosis upon **C:** bacterial treatment and **D:** TLR stimulation. **E:** monocyte CD52-expression (%) of previously defined CD52^high^ and CD52^low^ samples after migration. Phagocytosis capacity (% and MFI) of **F:** HC monocytes and **G:** previously defined CD52^high^ cirrhotic monocytes upon plasma conditioning from different disease stages. PLC plasma levels (ng/ml) **H:** at different stages of cirrhosis and **I:** correlate negatively with monocyte CD52-expression and positively with sCD52 plasma levels in cirrhosis patients. PLC plasma levels (ng/ml) **J:** related to survival, infection, HE, ascites and cirrhosis aetiology and **K**: predicts survival and infections. **L:** CD52-expression (% and MFI) of monocytes at different disease stages upon treatment with recombinant PLC. *p<0.05/**p<0.01/***p<0.001/****p<0.0001 (Friedman-, Kruskal-Wallis-, Wilcoxon-, Mann-Whitney test)

### PLC downregulates CD52 on monocytes and indicates infection and low transplant-free survival

PLC is known to shed GPI-anchored proteins [17], hence represents a putative candidate to downregulate surface CD52. Plasma levels of PLC were below detection limit in compensated and NAD cirrhosis, but significantly increased in AD/ACLF (Fig. 7H). In patients PLC levels correlated negatively with CD52-expression on monocytes and positively with sCD52 levels in the plasma (Fig. 7I). Plasma PLC elevation was clinically significant, as it was associated with low TFS. Patients with high PLC plasma levels were more likely to develop an infection within 4 weeks after recruitment. Furthermore, high PLC levels were associated with development of HE and ascites, but independent of the underlying aetiology (Fig. 7J). PLC levels predicted 1-year TFS (AUROC 0.7880) and the development of infection (AUROC 0.7269) (Fig. 7K).

To investigate whether PLC cleaves or downregulates CD52-expression on monocytes *ex vivo*, we incubated monocytes from HC and cirrhosis patients at different stages with PLC. We observed a significantly lower expression of CD52 on monocytes from compensated and NAD cirrhosis patients after PLC treatment, while no significant change was detected on CD52^low^ monocytes from HC and AD/ACLF patients (Fig. 7L).

### Physiological CD52 expression on liver macrophages is reduced in liver cirrhosis

In relation to the activated macrophage-like state of circulating CD52^high^ monocytes, liver macrophages isolated from resection tissue of patients with compensated cirrhosis and histologically normal liver were investigated. Human hepatic macrophages highly expressed CD52 in the physiological state. However, in cirrhosis, CD52-expression was reduced (Fig. 8A). Immunophenotyping of CD52^high^ hepatic macrophages revealed CD68^+^CD14^+^CD16^high^HLA-DR^high^ cells similar to CD52^high^ monocytes (Fig. 8B, C). CD52^high^ hepatic macrophages expressed higher levels of CD206 and AXL (Fig. 8C). Hepatic macrophages showed lower phagocytosis capacity in cirrhotic tissues, while CD52^high^ macrophages showed higher phagocytosis capacity compared to CD52^low^ similar to CD52^high^ monocytes in the circulation (Fig. 8D, E).

**Figure 8:**
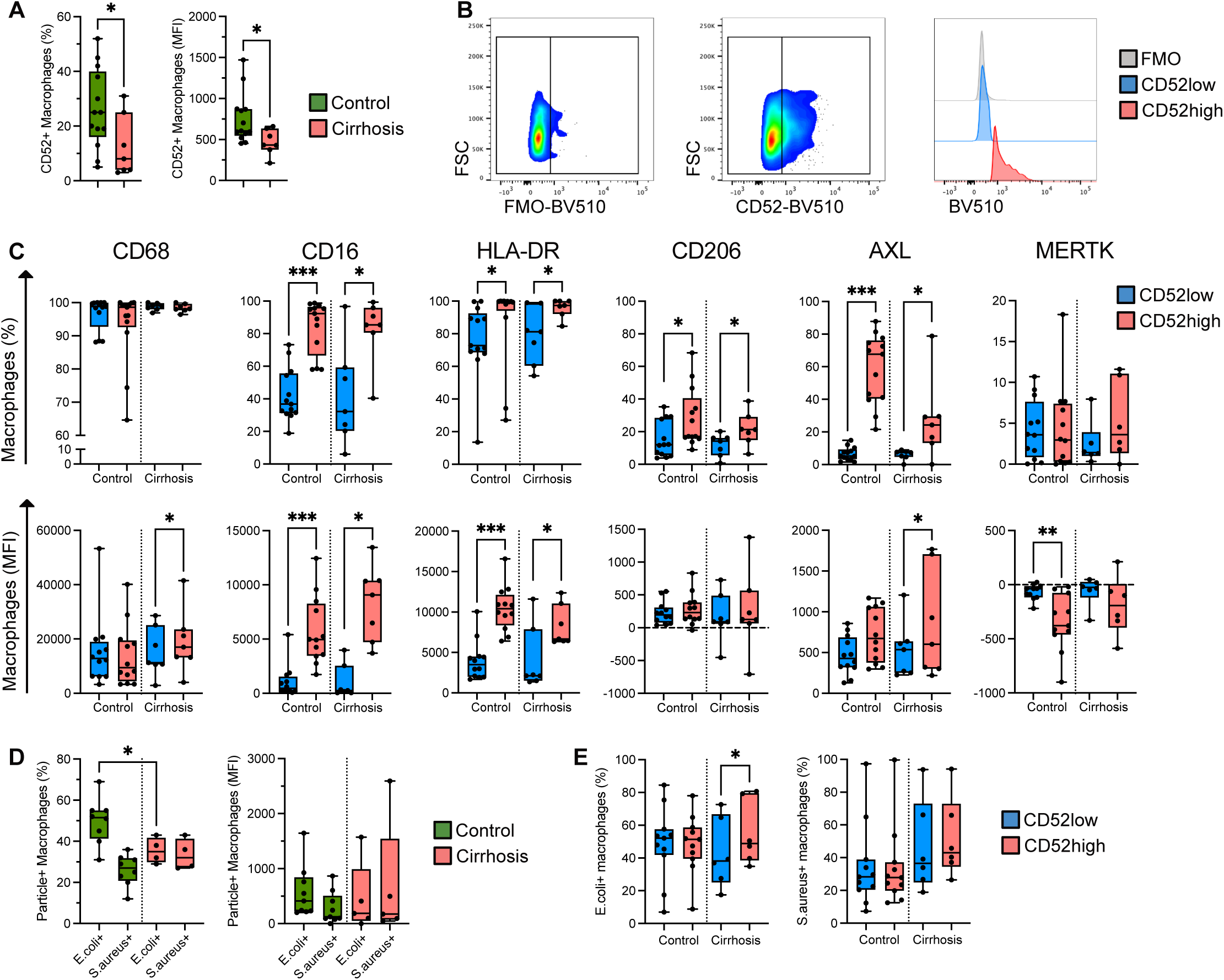
CD52 expression on isolated human hepatic macrophages *ex vivo*. **A:** CD52 expression (% and MFI) on human hepatic macrophages. **B:** Gating strategy and representative histograms to identify CD52^high^ and CD52^low^ macrophages. Forward scatter (FSC), fluorescence minus one (FMO). **C:** Immunophenotyping of CD52^high^ and CD52^low^ macrophages in cirrhosis. Phagocytosis capacity (% and MFI) of macrophages in **D:** control vs. cirrhosis and **E:**CD52^high^ vs. CD52^low^ macrophages. *p<0.05/**p<0.01/***p<0.001/****p<0.0001 (Mann-Whitney-, Wilcoxon-, Kruskal-Wallis-test)

## Discussion

The work delineated in this manuscript transcriptionally and systematically dissects monocyte states circulating in patients with compensated and decompensated cirrhosis, hereby recapitulating the reduction of non-classical monocytes and the accumulation of M-MDSC with disease severity, and identifying previously unknown distinct changes within in the classical monocyte subset involving upregulation of genes involved in resolution of inflammation and macrophage differentiation. We specifically observed that monocytes from patients with compensated disease or NAD over-expressed CD52, relating to an activated monocyte state with enhanced innate immune function, which was associated with survival. Cleavage or inhibition of CD52 by contrast as presented in the stage of AD/ACLF abrogated immune responses and implied poor prognosis, the process may be mediated by PLC. Stabilisation of CD52 on monocytes therefore represents a potential immunotherapeutic strategy worth exploring.

Cirrhosis associated immune dysfunction involves changes in the prevalence of certain circulating monocyte states, including a reduction in non-classical monocytes that are recruited to the injured liver [6]. Our scRNA-seq data clearly confirmed this finding, distinctly identifying non-classical, intermediate and classical monocytes by their transcriptional profile and showing a reduction of the circulating non-classical subset in the NAD stage. Additionally, our dataset confirms the accumulation of M-MDSC in the NAD stage, as previously described [7, 8, 29, 30]. Although this scRNA-seq study did not include samples from AD/ACLF patients, that previously showed highest circulating numbers of the immunosuppressive M-MDSC [7]. Given the high cellular plasticity [8, 31] and reversibility [23], it is essential to thoroughly dissect the clinical conditions and stages of cirrhosis in translational studies.

Given other distinct circulating monocyte states have been reported to occur at certain stages of cirrhosis, e.g. monocytes expressing TAM receptors MERTK or AXL [6, 8, 23], our scRNA-seq data revealed more monocytic states within the classical subset which differed between HC compensated and NAD cirrhosis, respectively. While in health cluster 6 characterised by genes required for cell degradation and recycling predominated the classical monocytes, we observed a shift towards cluster 5 enriched in genes relevant for phagocytosis in compensated cirrhosis and towards cluster 7 (M-MDSC) in NAD, as discussed. The data may imply activation of survival signals in monocytes in cirrhosis, confirming that viability of monocytes was not reduced even in AD/ACLF [7]. Also, it highlights the importance of phagocytosis in circulating monocytes in the context of pathological bacterial translocation [7, 32].

As previously described, we found the TAM receptor *MERTK,* important for the inhibition of TLR signalling cascades and efferocytosis in the context of resolution of inflammation [22, 33], upregulated on monocytes from patients with cirrhosis, again highlighting the relevance of *MERTK* as a biomarker of immuneparesis, and potential immunotherapeutic target worth evaluating [6]. Also *Gas6*, ligand and activator of the TAM receptors MERTK and AXL, was upregulated on CD14+ monocytes in cirrhosis. This may relate to the upregulation of *MERTK* or AXL on monocytes [8], of note the source of *Gas6* is not restricted to monocytes and macrophages, RNA and protein are expressed in several tissues and secreted to the blood. We previously observed a significant upregulation of GAS6 in hepatic stellate cells from cirrhosis patients [23].

Upregulation of inflammatory pathways involving high levels of cytokines in patients with cirrhosis was repeatedly reported previously and led to the systemic inflammation hypothesis [1, 2]. Due to the transition of stages during cirrhosis progression and inconsistencies in the nomenclature of stages over the years [34], the dynamics of inflammatory responses across stages remains insufficiently understood. The upregulation of genes involved in inflammatory pathways including humoral responses but also interferon signalling in monocytes from patients with cirrhosis confirms previous observations, while the lower expression in NAD compared to compensated cirrhosis is novel and confirms the concept that attenuated inflammatory responses may occur in NAD and prevail in AD/ACLF as previously noted [6, 7]. In this context the upregulation of genes involved in phagocytosis, plasma membrane organisation and endocytosis in both compensated and NAD stages is also highly interesting. We had previously reported, that phagocytosis was intact, highly active in cirrhosis, unless AD/ACLF occurred [7, 8], and indeed in this manuscript we revealed, that CD52^high^ monocytes, enhanced in compensation and NAD, were active phagocytes maintaining the patients’ infection defence. Similar findings have recently been reported from scRNA-seq/snRNA-seq from hepatic monocytes derived from cirrhotic livers [35].

Deterioration of synthetic liver function, e.g. protein synthesis, in cirrhosis is known, but its onset in relation to the dynamic decompensation stages remains poorly understood, especially the presence of coagulopathy and its clinical implications is an important focus of discussion and research [36]. It is interesting to note, that genes and cascades involved in coagulation were upregulated in monocytes from cirrhosis patients, also in the NAD stage, supporting enhanced rather then failed coagulation as conceptualised recently [37]. Complement pathways in cirrhosis were significantly less well studied [38], however in this dataset, complement genes and pathway were clearly enhanced in cirrhosis, even in the NAD, indicating preservation of soluble components of the immune system.

The upregulation of CD52 as well as PLC and other phosphatidyl signalling pathway components on monocytes from patients with liver cirrhosis in compensated and NAD stages is novel and highly interesting in relation to the heretofore poorly understood function of CD52 on monocytes on the one hand, and the beneficial prognostic profile on the other, highlighting its potential as future therapeutic target overcoming immuneparesis in cirrhosis. In healthy conditions CD52-expression is highest on B and T cells, while among myeloid cells CD16+ monocytes and mDC express the highest CD52 levels [39]. CD52 on T cells limits their activation [9] and sCD52 inhibits TLR activation of NF-kB and suppresses inflammation in THP-1 cells [11].

In our cohort of cirrhosis patients in different disease stages, we clearly observed, that CD14+ monocytes upregulated the expression of CD52 during evolution of cirrhosis, in stages with acceptable morbidity, immune control and prognosis. CD52^high^ cells were characterised as a CD14^+^CD16^high^HLA-DR^high^ mature monocytes with high phagocytosis capabilities, required for pathogens clearance and initiation of immune responses, high cytokine responses enhanced upon TLR challenge, and high potential to adhere to and migrate across endothelial barriers, as required for the recruitment of monocytes to inflamed tissues, however low potential to activate T cell proliferation. Enhancing these selective monocyte functions appear beneficial in the context of emerging pathological bacterial translocation [40] and the sporadic occurrence of sterile or infectious inflammation of the liver and other tissues [4] and may therefore be seen as a rescue process enhancing innate immune function in the condition of cirrhosis with emerging architectural changes of the liver sinusoids and its filter function between portal and systemic circulation. This may explain the clear association of CD52^high^ monocyte prevalence with survival in our cohort.

The question emerges which signals lead to the upregulation of CD52-expression as early as cirrhosis emerges. *In vitro* we revealed, that plasma components of cirrhosis patients lead to the upregulation of CD52 on healthy monocytes, similarly upregulation was also observed after phagocytosis of bacterial particles, whereas stimulation with TLR ligands such as LPS was insufficient. Another process upregulating CD52, was migration across liver sinusoidal endothelia. These findings implicate that the enhanced expression of CD52 may originate from different processes ongoing during cirrhosis evolution – including the presence of circulating bacterial products, other changes in the plasma composition, and migration of monocytes towards tissues and potentially back to the circulation.

By contrast, in stages of AD/ACLF CD52^high^ monocytes disappeared from the circulation, which was associated with poor survival. CD52^low^ monocytes had reduced phagocytosis potential, low cytokine responses, and low migratory potential. A similar immunosuppressive profile occurred following treatment with alemtuzumab, but also PLC treatment. PLC has been shown to cleave CD52 on T cells, and release sCD52 to the plasma [17]. Interestingly our cohort of patients, simultaneously revealed CD52^low^ monocytes and high plasma levels of both PLC and sCD52. The source of PLC in the plasma remains unknown, there are six different isoforms and various splicing variants in humans, which are expressed in diverse tissues [41]. In the context of liver cirrhosis phospholipase A release from liver and kidney has been reported in rats [42]. The increased release of PLC in AD/ACLF has not been reported; it may origin from sources of tissue inflammation precipitating AD, e.g. the peritoneum or kidney, less likely the inflamed liver itself since it is implicated in the pathogenesis of numerous inflammatory diseases [43].

CD52 has been described as a marker associated with survival also in another context. A study evaluating dynamical changes in gene expression on PBMC by single scRNA-seq over 6 hours in patients with gram negative sepsis revealed that CD52-expression increased over 6 h in T and B cells of survivors, but not in sepsis non-survivors, and identifying CD52 as a biomarker correlating with beneficial outcome [13]. Moreover, CD52 overexpression in breast cancer tissue correlated positively with M1-macrophage- and negatively with M2-macrophage infiltration and also survival in [44].

In this paper we also provide evidence, that the distinct functional phenotype of CD52^high^ monocytes can be recapitulated *in vitro* when overexpressing CD52 with a viral vector construct. The experiments give rise to the question, whether in the future, stabilisation of CD52-expression on immune cells could be used for therapeutic immune-modulation? In the endoplasmic reticulum GPI-anchored proteins are stabilized by p24 complex during posttranslational modification and export [45]. If stabilisation of CD52 similar to overexpression could enhance immune responses such as phagocytosis, cytokine responses and migration across endothelia in cirrhosis patients, this may lower their susceptibility to infection and prevent infectious episodes, AD and death – allowing for the liver to regenerate or bridge to transplantation.

In summary, we detail circulating monocytes by transcriptome analysis in patients with compensated and NAD cirrhosis, recapitulating the loss of CD14+CD16++ monocytes and the emergence of M-MDSC in decompensated stages. Moreover, we detect CD14++ monocytes with a signature of enhanced phagocytosis emerging early in cirrhosis in adaptation to the changes in the hepatic architecture and composition of portal and systemic milieu. For the first time, we observe the upregulation of CD52 on functionally activated monocytes and in correlation with survival. In AD/ACLF by contrast, these activated CD52^high^ monocytes were lost, by cleavage from PLC potentially released from sites of inflammation, relating to poor prognosis. Regulation of CD52 on monocytes may be a mechanism for future exploration as immunotherapy in patients with cirrhosis.

## Data availability statement

scRNA-seq processed data (UMI count table) will be made available on GEO (submission ongoing). Raw sequencing data are considered sensitive and thus cannot provided on public data repositories. Upon request to the authors, a data access agreement can be signed between involved institutions to share these raw data on a case-by-case basis.

## Acknowledgments

The authors thank all patients who consented to participate in this study and all staff at the University Hospital Basel and the Cantonal Hospital St.Gallen involved in these patients’ care. The authors also thank the Medical Research Center of the Cantonal Hospital St.Gallen and the Department of Biomedicine of the University of Basel for infrastructural support. In particular, the authors are grateful to the Gastroenterology Research Group, led by Prof. J. Niess, Department of Biomedicine, for methodological and conceptual advice. The authors also thank the core facilities including the bioinformatics Core and Flow cytometry core facilities. The authors thank Deniz Kaymak (Prof. G. Hutter, Brain Tumor Immunotherapy and Biology Research Group, Department of Biomedicine) for the kind gift of packaging plasmids psPAX2 and pMD2.G and methodological support.

Calculations were performed at sciCORE (http://scicore.unibas.ch/) scientific computing center at the University of Basel. Graphical abstract was created with BioRender.com

## Author contributions

study concept and design: CB, RGB, BL

acquisition of data: AG, RGB, JR, ML, H-WC, EEF, M-AM, OTP, PK-H, MM, SS, DS, MH, CW, BL, CB

analysis and interpretation of data: AG, RGB, JR, ML, H-WC, EEF, M-AM, OTP, CB

drafting of the manuscript: AG, CB

critical revision of the manuscript for important intellectual content: RGB, JR, ML, H-WC, EEF, M-AM, OTP, PK-H, MM, SS, DS, MH, CW, BL

statistical analysis: AG, RGB, JR, ML

obtained funding: CB

administrative, technical, or material support: H-WC, M-AM, PK-H, MM, SS, DS, MH, CW, BL

study supervision: CB

