## Supplementary Material for "Distinct circulating monocytes up-regulate CD52 and sustain innate immune function in patients with cirrhosis unless acute decompensation emerges"

### Methods

#### Clinical, Haematologic, and Biochemical Parameters

Episodes of infection, disease severity scores (Child Pugh, Model of end stage liver disease [MELD] [1], Chronic liver failure-acute decompensation [CLIF-AD] [2]) were calculated, and stages of AD/ACLF as defined by the CLIF-SOFA classification [3] were documented.

#### Single cell RNA sequencing methods

STARsolo [4] (version 2.7.10a) was used to perform sample and cell demultiplexing, and alignment of reads to the human genome (hg38) and UMI counting on gene models from Ensembl 104 excluding Locus Reference Genomic sequences (with options --outFilterType=BySJout --outFilterMultimapNmax=10 --outSAMmultNmax=1 --outFilterScoreMin=30 --soloCBmatchWLtype=1MM_multi_Nbase_pseudocounts --soloUMIfiltering=MultiGeneUMI_CR --soloUMIdedup=1MM_CR --soloType=CB_UMI_Simple --soloStrand=Forward --clipAdapterType=CellRanger4 --soloUMIlen=12 --soloCellFilter=None --soloMultiMappers=EM).

Processing of the UMI counts matrix was performed using Bioconductor (version 3.16) packages, notably DropletUtils [5] (function *emptyDrops* using 5000 iterations, the option test.ambient=TRUE, a lower threshold of 100 UMIs and an FDR threshold of 0.1%), scran [6, 7] and scater [8], following mostly the steps illustrated in the OSCA book [9] (http://bioconductor.org/books/3.16/OSCA/). Cells with 0% or more than 12% of UMI counts attributed to mitochondrial genes, with less than 3000 UMI counts, or with less than 1500 detected genes were excluded. The presence of doublet cells was investigated with the scDblFinder package [10] (version 1.12.0), and suspicious cells were filtered out (score<0.2). The package SingleR [11] (version 2.0.0) was used for cell-type annotation of the cells using the following celldex human references: MonacoImmuneData, HumanPrimaryCellAtlasData, BlueprintEncodeData and DatabaseImmuneCellExpressionData. The assigned pruned labels were used and the cells not annotated to monocyte subtypes in these references were excluded. Additionally, cells from one outlier patient (with compensated cirrhosis) were excluded because unlike any of the other patients, it displayed a widespread increased interferon response.

Systematic differences between samples were removed using the fastMNN function (k=30) of the batchelor package [12] (version 1.14.1). A UMAP embedding for visualization of cells was calculated using the batch-corrected low-dimensional coordinates for cells (n_neighbors=50, min_dist=0.5). Shared nearest-neighbor graph clustering was performed using the Leiden algorithm for finding community structure (k=10), resulting in 17 clusters which were merged into 7 clusters based on manual inspection of the cluster-specific genes, to reflect the continuum between classical monocytes, non-classical monocytes and M-MDSCs.

Differences in relative abundance of clusters across patient groups were tested using limma-voom [13]. This method was shown to be able to handle well the overdispersion in the cell type proportions estimates typically observed with single cell technologies [14]. Differential abundance of cell types was considered to be significant at a false discovery rate (FDR) lower than 10%.

Differential expression across patient groups was tested using a pseudo-bulk approach, summing the UMI counts of cells from each cluster in each sample. The aggregated samples were then treated as bulk RNA-seq samples [15]. The package edgeR [16] (version 3.40.2) was used to perform TMM normalization [17] and to test for differential expression (*glmLRT*). Genes with an FDR lower than 10% were considered differentially expressed. Gene set enrichment analysis was performed with the function camera [18] on gene sets from the Molecular Signature Database (MSigDB) collection, notably the Hallmark collection [19, 20] (version 7.5.1). We retained only sets containing more than 5 genes, and gene sets with an FDR lower than 10% were considered as significant.

#### Monocyte and T cell Isolation

Peripheral blood mononuclear cells (PBMCs) were isolated from heparinised whole blood samples and monocytes and T cells were isolated by magnetic-activate cell sorting (MACS) using Pan Monocyte Isolation Kit (Miltenyi Biotec 130-096-537) or Pan T Cell Isolation Kit (Miltenyi Biotec 130-096-535) according to the manufacturers protocol.

#### Human Tissue Macrophage Isolation

In sum, small resection samples were mechanically disrupted using a filter mesh, whereas large resection samples were processed using a Stomacher 400 (Seward).

#### Flow Cytometry based Phenotyping

At least 100µl whole blood or 100.000 cells, respectively, were incubated in 100µl FACS buffer. Cells were fixed, permeabilised using the Foxp3 / Transcription Factor Staining Buffer Set (ThermoFisher SCIENTIFIC, Cat.: 00-5523-00)) and stained with antibodies for flow cytometry (Table S1). Flow cytometry data was analysed using FlowJo software (V.10.8.1, Becton Dickinson & Company) as previously described [21].

#### Flow Cytometry based Phagocytosis Assay

Cells were incubated with pHrodo *E.coli* ( ThermoFisher SCIENTIFIC, Cat.: P35361) or *S.aureus* (ThermoFisher SCIENTIFIC, Cat.: A10010) Red BioParticles according to the manufacturer's protocol. At least 100µl whole blood or 100.000 cells in 100µl FACS buffer supplemented with 10% human AB serum were incubated with 5 µl BioParticles for 30 min at 37°C. Cells were fixed and stained with antibodies for flow cytometry as previously described [22].

#### Assessment of Intracellular Cytokine Responses to TLR Stimulation

Monocytes were stimulated with Lipopolysaccharide (LPS), synthetic triacylated lipopeptide (Pam3CSK4) for 5h. BD GolgiStop$^{TM}$ Protein Transport Inhibitor (BD Biosciences, Cat.: 554724) was added after 1h to accumulate cytokines in the Golgi complex for 4h. Monocytes were fixed and stained intracellularly for flow cytometry.

#### Mixed Lymphocyte Reaction

Monocytes from HC and monocytes from patients with cirrhosis were co-cultured with T cells from a healthy donor in a 1:1 ratio. T cell proliferation was induced with anti-CD2/CD3/CD28 beads (T cell Activation/Expansion Kit, Miltenyi Biotec, 130-091-441). T cells were stained with carboxyfluorescein succinimidyl ester (CFSE) at day 0. Proliferation was assessed after 24h of co-culture by flow cytometry.

#### Adhesion Assay in Static Conditions to Immobilized ICAM1-Fc, ICAM2-Fc and VCAM1-Fc

In short, 96-well clear bottom, black plates were coated with 10 µg/mL ICAM1-Fc (BioLegend, Cat.: 552906), ICAM2-Fc (BioLegend, Cat.: 768506) and VCAM1-Fc (BioLegend, Cat.: 553706) in PBS with $Ca^{2+}/Mg^{2+}$ (ThermoFisher SCIENTIFIC, Gibco$^{TM}$, Cat.: 14040133) for 2h in 37°C. Control wells were coated with 10 µg/ml Bovine Serum Albumin (BSA) (CARL ROTH). Plates were washed with PBS and monocytes were stained with CFSE. 1 x 10^5 monocytes were plated onto each well and allowed to adhere for 20 min. Fluorescence of loaded cells was measured on a plate reader. Plates were further washed with PBS with $Ca^{2+}/Mg^{2+}$ and fluorescence of adherent cells was measured. The coefficient of adhesion was calculated:

$$MFI\ ratio = \frac{\text{fluorescence of adeherent cells (MFI)}}{\text{fluorescence of loaded cells (MFI)}}$$

#### In Vitro Inhibition of CD52

Isolated monocytes were incubated with alemtuzumab as previously described [23]. 20 µg/ml Alemtuzumab (biotechne, Human CD52 research grade Alemtuzumab Biosimilar Antibody, Cat.: MAB9889) for 1h at 37°C. Unbound monoclonal antibody (mAB) was washed off before proceeding further. Immunophenotyping, phagocytosis capacity, cytokine production, adhesion-, migration- and T cell activation potential were assessed.

#### Generation of THP-1 Cell Line Stably Expressing CD52

THP-1 cells were transduced pLV[Exp]-CMV>hCD52[NM_001803.3] vector and THP-1-CD52+ cells were sorted by flow cytometry. THP-1 and THP-1-CD52+ cell lines were cultured in Rosewell Park Memorial Institute 1640 medium (RPMI 1640, Sigma-Aldrich, Cat.: R8758) supplemented with 10% heat-inactivated fetal bovine serum (FBS, ThermoFisher SCIENTIFIC, Gibco$^{TM}$, Cat.: 10500064) and Penicillin-Streptomycin-Glutamine (Pen/Strep 100X, ThermoFisher SCIENTIFIC, Gibco$^{TM}$, Cat.: 10378016).

#### Soluble CD52

and according to the manufacturers protocol (CUSABIO® Human CAMPATH-1 antigen (CD52) ELISA kit, Cat.: CSB-EL004943HU).

#### Conditioning with Plasma

Monocytes were from HC expressing low CD52 levels (Median: 32.4%, 348.5MFI) (<40% / <600MFI CD52-expressing monocytes) were incubated for 18h in RPMI1640 with 25% plasma from HC, patients with cirrhosis or AD/ACLF. Monocytes from patients with cirrhosis expressing high CD52 (Median: 75.80% / 72.65%, 644MFI / 496MFI) (>60% / >600MFI CD52-expressing monocytes) were incubated for 18h in RPMI1640 with 25% plasma from HC or patients with AD/ACLF [22]. Immunophenotyping and phagocytosis capacity was assessed.

### Supplementary Table 1: Cirrhosis Patients

**Supplementary Table 1: Clinical characteristics of patients with cirrhosis compared to healthy controls**

| Variables, median (range) | Cirrhosis (n = 63) | Healthy controls (n = 27) |
|---|---|---|
| **Age (y)** | 62 (21 - 85)*** | 42 (26 - 65) |
| **Sex (m : f)** | 40 : 23 | 8 : 19 |
| **Underlying liver disease(s)** | | |
| ALD | 40 | NA |
| NAFLD | 21 | NA |
| HCV | 7 | NA |
| HBV | 5 | NA |
| HBV/HDV Superinfection | 1 | NA |
| HEV | 1 | NA |
| PBC | 1 | NA |
| AIH | 1 | NA |
| Drug Toxic | 1 | NA |
| unknown | 2 | NA |
| **Death (1 year)** [%] | 9 [14] | - |
| **Child Pugh** | 7 (5 - 13) | NA |
| **Model for End-Stage Liver Disease (MELD)** | 14 (5 - 28) | NA |
| **BMI (kg/m$^2$)** | 27,4 (17,5 - 38,8)*** | 22.3 (18,6 - 27,9) |
| **$Na^+$ (mmol/L)** | 139 (124 - 143) | 139,5 (138 - 141) |
| **$K^+$ (mmol/L)** | 4,1 (2,9 - 4,9) | 4 (3,8 - 4,3) |
| **Creatinine (µmol/L)** | 72 (36 - 320) | 74 (58 - 80) |
| **Urea (mmol/L)** | 5,1 (1,5 - 20.9) | 5,2 (3,4 - 5,8) |
| **Bilirubin (µmol/L)** | 23,6 (2,6 - 644,5)* | 9,1 (7 - 11,9) |
| **Aspartat-Aminotransferase (AST)** [U/L] | 45 (12 - 304)** | 21 (20 - 28) |
| **Alanin-Aminotransferase (ALT)** [U/L] | 36 (10 - 153) | 21 (20 - 28) |
| **Gamma-Glutamyltransferase (GGT)** [U/L] | 79 (11 - 1760)* | 23,5 (13 - 83) |
| **Alkaline-Phosphatase (AP)** [U/L] | 113 (50 - 572)*** | 55,5 (38 - 60) |
| **International Normalised Ratio (INR)** | 1,2 (0,9 - 2,7)* | 1 (1 - 1,2) |
| **Albumin (g/L)** | 33 (17 - 46) | 39 (38 - 40) |
| **Hemoglobin (g/L)** | 122 (68 - 165) | 136 (132 - 171) |
| **Hematocrit (%)** | 0.37 (0.14 - 0.5) | 0,4 (0,39 - 0,51) |
| **Leukocytes (G/L)** | 5.2 (2.3 - 12-7) | 6 (4.2 - 9.1) |
| **Neutrophils (G/L)** | 3.9 (0.7 - 10.1) | 3.2 (2.4 - 5.8) |
| **Eosinophils (G/L)** | 0.13 (0.01 - 0.84) | 0.06 (0.05 - 0.35) |
| **Basophils (G/L)** | 0.03 (0.01 - 0.27) | 0.04 (0.01 - 0.06) |
| **Lymphocytes (G/L)** | 1.08 (0.23 - 3) | 2.05 (1.11 - 3.08) |
| **Monocytes (G/L)** | 0.37 (0.04 - 1.06) | 0.37 (0.18 - 0.59) |
| **Platelets (G/L)** | 106 (21 - 675) | 250 (165 - 379) |
| **C-Reactive Protein (CRP)** [mg/L] | 6.8 (0,3 - 54) | 0.35 (0.3 - 0.8) |

ALD, alcoholic liver disease; NAFLD, Non-alcoholic fatty liver disease; HCV, hepatitis C virus; HBV, hepatitis B virus; HDV, hepatitis D virus; HEV, hepatitis E virus; PBC, primary biliary cirrhosis; AIH, autoimmune hepatitis; BMI, body mass index; Data indicated as median with (minimum and maximum) and [percentage]. *$P$=0.05, **$P$=0.01, ***$P$=0.001 indicate cirrhosis vs healthy controls, comparisons by Mann-Whitney *U* Tests.

### Supplementary Table 2: Liver Resection Patients

**Supplementary Table 2: Clinical characteristics of patients with cirrhosis compared to histologically normal controls**

| Variables, median (range) | Cirrhosis (n = 12) | Controls (n = 16) |
|---|---|---|
| **Age (y)** | 62 (48 – 84) | 62.5 (28 – 78) |
| **Sex (m : f)** | 7 : 5 | 5 : 11 |
| **Underlying liver disease(s)** | | |
| ALD | 9 | - |
| NAFLD | 2 | - |
| HCV | 3 | - |
| Echinococcus | - | 3 |
| Liver Metastasis | - | 13 |
| **Death (1 year)** [%] | | |
| **Child Pugh** | 5 (5 – 9) | NA |
| **Model for End-Stage Liver Disease (MELD)** | 8 (6 – 15) | NA |
| **$Na^+$ (mmol/L)** | 139 (132 – 143) | 140 (136 – 141) |
| **$K^+$ (mmol/L)** | 4 (3.7 – 4.7) | 4.2 (3.8 – 4.5) |
| **$Ca^{2+}$ (mmol/L)** | 2.4 (2.1 – 2.6) | 2.4 (2.2 – 2.5) |
| **Creatinine (µmol/L)** | 60 (48 – 132) | 60 (42 – 100) |
| **Glomerular Filtration Rate (GFR)** [**ml/min/1.7**] | 94 (51 – 109) | 96 (62 – 127) |
| **Urea (mmol/L)** | 5.3 (2.6 – 10.7) | 5 (3.8 – 8.6) |
| **Uric Acid (µmon/L)** | 276.5 (149 – 463) | 226 (2.54 – 546) |
| **Bilirubin (µmol/L)** | 10.8 (4.6 – 22.1)** | 5.1 (2.5 – 14.2) |
| **Aspartat-Aminotransferase (AST)** [**U/L**] | 33 (17 – 83) | 29 (18 – 52) |
| **Alanin-Aminotransferase (ALT)** [**U/L**] | 26 (13 – 48) | 28 (14 – 56) |
| **Gamma-Glutamyltransferase (GGT)** [**U/L**] | 57 (12 – 796) | 61 (16 – 858) |
| **Protein Total (g/L)** | 71 (62 – 78) | 73 (66 – 81) |
| **Alkaline-Phosphatase (AP)** [**U/L**] | 80 (44 – 557) | 105 (44 – 487) |
| **Lactate Dehydrogenase (LDH)** [**U/L**] | 209 (134 – 289) | 231 (179 – 513) |
| **International Normalised Ratio (INR)** | 1.1 (1 – 1.9)** | 1 (0.9 – 1.1) |
| **Albumin (g/L)** | 38 (25 – 43) | 37 (29 – 42) |
| **Hemoglobin (g/L)** | 137 (88 – 150) | 121 (104 – 155) |
| **Hematocrit (%)** | 0.4 (0.28 – 0.43) | 0.36 (0.31 – 0.43) |
| **Leukocytes (G/L)** | 5.24 (2.74 – 8.67) | 5.58 (3.13 – 12.02) |
| **Erythrocytes (G/L)** | 4.31 (3.03 – 5.3) | 4.14 (3.64 – 5.01) |
| **Neutrophils (G/L)** | 3.14 (1.97 – 6.23) | 3.17 (2.02 – 4.91) |
| **Eosinophils (G/L)** | 0.07 (0.02 – 0.16) | 0.12 (0.03 – 0.59) |
| **Basophils (G/L)** | 0.03 (0.01 – 0.03) | 0.02 (0.01 – 0.08) |
| **Lymphocytes (G/L)** | 0.96 (0.48 – 1.85) | 1.46 (1.05 – 2.37) |
| **Monocytes (G/L)** | 0.31 (0.19 – 0.61) | 0.32 (0.22 – 1) |
| **Platelets (G/L)** | 151 (99 – 279)* | 238 (100 – 363) |
| **C-Reactive Protein (CRP)** [**mg/L**] | 2.6 (0.3 – 74.2) | 2.1 (0.5 – 13.2) |

ALD, alcoholic liver disease; NAFLD, Non-alcoholic fatty liver disease; HCV, hepatitis C virus; PBC, primary biliary cirrhosis; AIH, autoimmune hepatitis; Data indicated as median with (minimum and maximum) and [percentage]. **P=0.05*, ***P=0.01*, ****P=0.001* indicate cirrhosis vs control, comparisons by Mann-Whitney *U* Tests.

### Supplementary Table 3: Reagents

**Supplementary Table 3: Reagent table**

| Reagent | Supplier | Catalogue number |
|---|---|---|
| Anti-Amyloid Precursor Protein antibody (Y188) (PE) | abcam | ab208744 |
| Human Axl Alexa Fluor® 488-conjugated Antibody | Biotechne | FAB154G |
| Brilliant Violet 510™ anti-human CD192 (CCR2) Antibody | BioLegend | 357218 |
| Brilliant Violet 421™ anti-human CD195 (CCR5) Antibody | BioLegend | 359118 |
| Brilliant Violet 421™ anti-human CD197 (CCR7) Antibody | BioLegend | 353208 |
| APC/Fire™ 750 anti-human CD3 Antibody | BioLegend | 300470 |
| BD Pharmingen™ PE Mouse Anti-Human CD11b | BD Biosciences | 555388 |
| BD Pharmingen™ PE-Cy7 Mouse Anti-Human CD14 | BD Biosciences | 557742 |
| PerCP/Cyanine5.5 anti-human CD14 Antibody | BioLegend | 325621 |
| APC anti-human CD15 (SSEA-1) Antibody | BioLegend | 301908 |
| Brilliant Violet 605™ anti-human CD15 (SSEA-1) Antibody | BioLegend | 323032 |
| BD Horizon™ BV650 Mouse Anti-Human CD16 | BD Biosciences | 563692 |
| BD Pharmingen™ APC Mouse Anti-Human CD19 | BD Biosciences | 555415 |
| APC/Fire™ 750 anti-human CD19 Antibody | BioLegend | 302258 |
| Brilliant Violet 650TM anti-human CD25 Antibody | BioLegend | 302634 |
| CD32 Monoclonal Antibody (6C4 (CD32)); FITC, eBioscience™ | ThermoFisher | 11-0329-42 |
| BD Horizon™ BV510 Mouse Anti-Human CD52 | BD Biosciences | 563305 |
| BD Pharmingen™ PE Mouse anti-Human CD52 | BD Biosciences | 562945 |
| APC/Fire™ 750 anti-human CD56 (NCAM) Antibody | BioLegend | 362554 |
| CD56 Antibody, anti-human, PE-Vio® 770, REAfinity™ | Miltenyi Biotec | 130-113-313 |
| BD Pharmingen™ FITC Mouse Anti-Human CD64 | BD Biosciences | 555527 |
| Brilliant Violet 785™ anti-human CD68 Antibody | BioLegend | 333826 |
| APC anti-human CD69 Antibody | BioLegend | 310910 |
| BD Pharmingen™ PE Mouse Anti-Human CD163 | BD Biosciences | 556018 |
| Brilliant Violet 421™ anti-human CD206 (MMR) Antibody | BioLegend | 321126 |
| Brilliant Violet 510™ anti-human CX3CR1 Antibody | BioLegend | 341622 |
| Brilliant Violet 510TM anti-human CD184 (CXCR4) Antibody | BioLegend | 306536 |
| PerCP/Cyanine5.5 anti-human HLA-DR Antibody | BioLegend | 307630 |
| BD Pharmingen™ FITC Mouse Anti-Human HLA-DR | BD Biosciences | 556643 |
| IL-10 Monoclonal Antibody (JES3-9D7), Alexa Fluor™ 488, eBioscience™ | ThermoFisher | 53-7108-42 |
| PE anti-human IL-6 Antibody | BioLegend | 501107 |
| Human IFN-alpha / beta R1 PE-conjugated Antibody | Biotechne | FAB245P |
| Human Mer APC-conjugated Antibody | Biotechne | FAB8912A |
| BD OptiBuild™ BV605 Mouse Anti-Human Mer | BD Biosciences | 748103 |
| BD Pharmingen™ Alexa Fluor® 647 Mouse Anti-Human MNDA | BD Biosciences | 566582 |
| PE anti-human Siglec-10 Antibody | BioLegend | 347604 |
| Human S100A8/S100A9 Heterodimer Alexa Fluor® 488-conjugated Antibody | Biotechne | IC9337G |
| PE anti-human Tim-4 Antibody | BioLegend | 354003 |
| APC anti-human CD284 (TLR4) Antibody | BioLegend | 312816 |
| Brilliant Violet 421™ anti-human TNF-α Antibody | BioLegend | 502932 |
| BD Horizon™ BV650 Mouse IgG1, k Isotype Control | BD Biosciences | 563231 |
| BD Pharmingen™ PE Mouse IgG1, κ Isotype Control | BD Biosciences | 555749 |
| BD Pharmingen™ FITC Mouse IgG1, κ Isotype Control | BD Biosciences | 555748 |
| APC Mouse IgG1, κ Isotype Ctrl Antibody | BioLegend | 400120 |
| APC/Fire™ 750 Mouse IgG1, κ Isotype Ctrl Antibody | BioLegend | 400196 |
| PerCP/Cyanine5.5 Mouse IgG2a, κ Isotype Ctrl Antibody | BioLegend | 400252 |
| BD Pharmingen™ PE-Cy™7 Mouse IgG2a, κ Isotype Control | BD Biosciences | 557907 |
| BD Pharmingen™ FITC Mouse IgG2a, κ Isotype Control | BD Biosciences | 555573 |
| Brilliant Violet 785™ Mouse IgG2b, κ Isotype Ctrl Antibody | BioLegend | 400355 |
| Mouse IgG1 Alexa Fluor® 488-conjugated Antibody | Biotechne | IC002G |
| Brilliant Violet 421™ Mouse IgG1, κ Isotype Ctrl Antibody | BioLegend | 400158 |
| CellTrace™ CFSE Cell Proliferation Kit, for flow cytometry | ThermoFisher | C34554 |
| eBioscience™ Fixable Viability Dye eFluor™ 455UV | ThermoFisher | 65-0868-18 |
| eBioscience™Foxp3 / Transcription Factor Staining Buffer Set | ThermoFisher | 00-5523-00 |
| Lysing Solution, Whole Blood Lysing Solution | ThermoFisher | GAS010 |
| pHrodo™ BioParticles™ Conjugates for Phagocytosis and Phagocytosis Kit, for Flow Cytometry | ThermoFisher | P35361 |
| pHrodo™ BioParticles™ Conjugates for Phagocytosis and Phagocytosis Kit, for Flow Cytometry | ThermoFisher | A10010 |

[19] Subramanian A, Tamayo P, Mootha VK, Mukherje S, Ebert BL, Gillette MA, et al. Gene set enrichment analysis: A knowledge-based

approach for interpreting genome-wide expression profiles. PNAS 2005;102.

[20] Liberzon A, Birger C, Thorvaldsdottir H, Ghandi M, Mesirov JP, Tamayo P. The Molecular Signatures Database (MSigDB) hallmark gene set collection. Cell Syst 2015;1:417-425.

[21] Abeles RD, McPhail MJ, Sowter D, Antoniades CG, Vergis N, Vijay GKM, et al. CD14, CD16 and HLA-DR reliably identifies human monocytes and their subsets in the context of pathologically reduced HLA-DR expression by CD14hi/CD16neg monocytes: Expansion of CD14hi/CD16pos and contraction of CD14lo/CD16pos monocytes in acute liver fail. Cytometry 2012;81A:823-834.

[22] Bernsmeier C, Triantafyllou E, Brenig R, Lebosse FJ, Singanayagam A, Patel VC, et al. $CD14^{+}$ $CD15^{-}$ HLA-$DR^{-}$ myeloid-derived suppressor cells impair antimicrobial responses in patients with acute-on-chronic liver failure. Gut 2018;67:1155-1167.

[23] Rudnik M, Rolski F, Jordan S, Mertelj T, Stellato M, Distler O, et al. Regulation of Monocyte Adhesion and Type I Interferon Signaling by CD52 in Patients With Systemic Sclerosis. Arthritis & Rheumatology 2021;73:1720-1730.

### Figure Legends

#### Supplementary Figure 1: Quality control of Single-cell data of human circulating monocytes

UMAP embedding showing the monocytes from the scRNA-seq dataset **A:** cells coloured according to patient samples, or **B** and **C:** disease stage, **D**: cells coloured according to label from unbiased annotation using Monaco et al. reference atlas, Cell Reports, 2019; cells coloured according to fraction of UMI counts mapped to genes from the mitochondrial genes or according to the number of detected genes. **E:** Cluster frequencies in absolute cell numbers and % of clusters and monocyte subsets separated by disease stage and each sample separately.

#### Supplementary Figure 2: DE genes in monocytes in cirrhosis

Heatmap of DE genes in monocytes comparing **A:** healthy vs. cirrhosis and **B:** compensated vs. NAD cirrhosis. The log-normalized expression levels were averaged across cells from the same cluster and sample, and averaged values were then centred and scaled per gene.

### Supplementary Figure 1

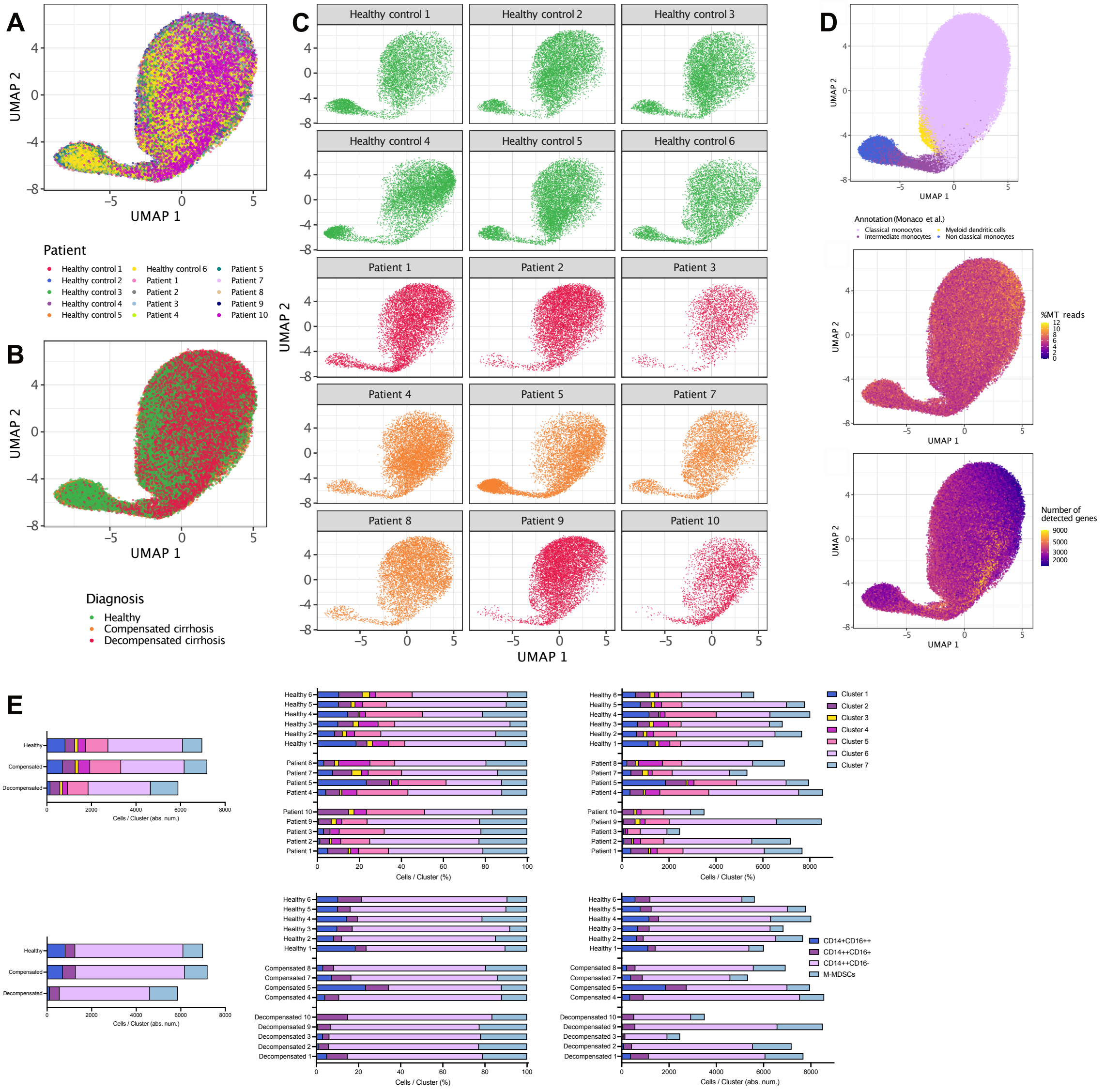

### Supplementary Figure 2

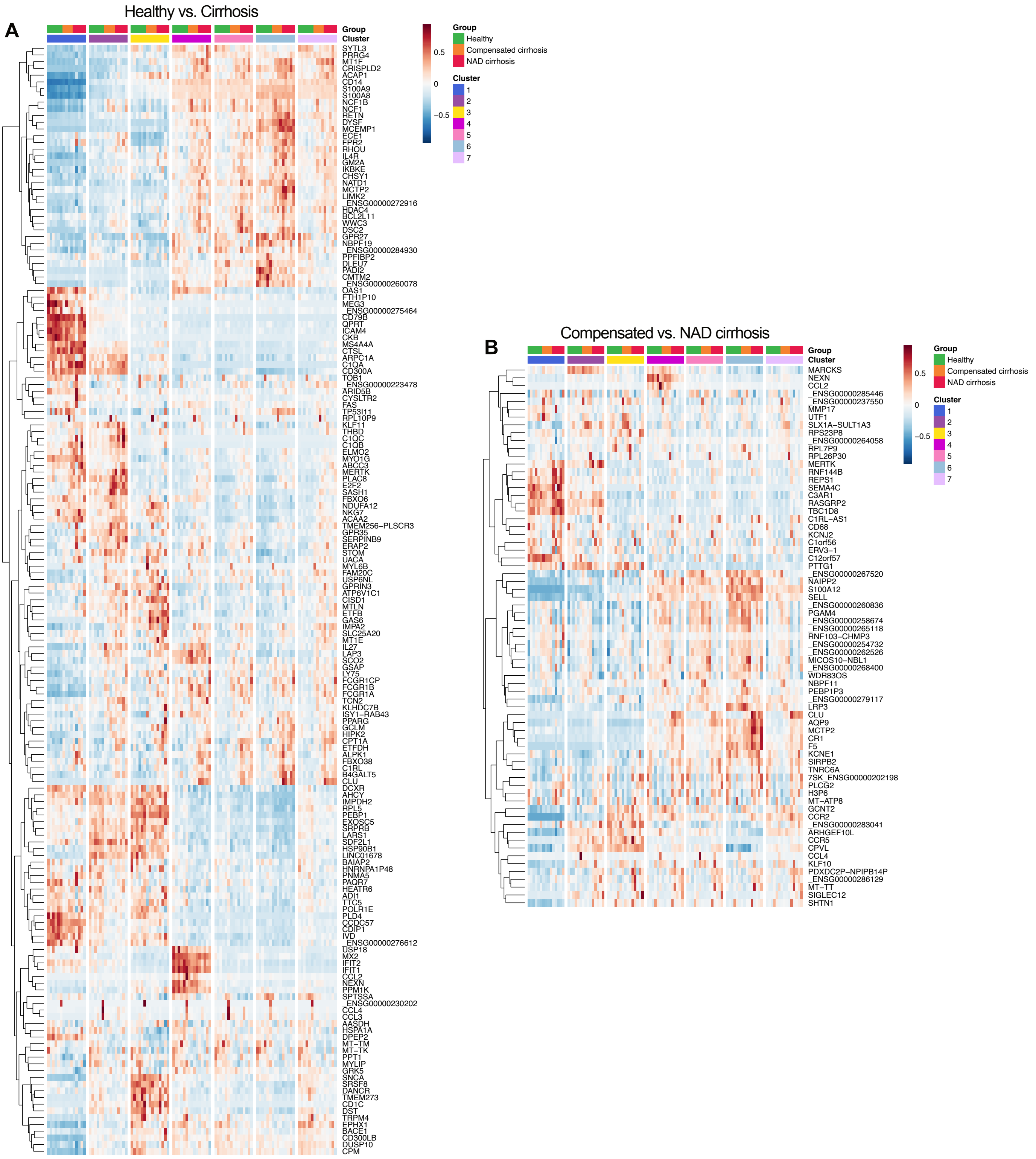
